# A dynamical model of the laminar BOLD response

**DOI:** 10.1101/609099

**Authors:** Martin Havlicek, Kamil Uludag

## Abstract

High-resolution functional magnetic resonance imaging (fMRI) using blood oxygenation dependent level-dependent (BOLD) signal is an increasingly popular tool to non-invasively examine neuronal processes at the mesoscopic level. However, as the BOLD signal stems from hemodynamic changes, its temporal and spatial properties do not match those of the underlying neuronal activity. In particular, the laminar BOLD response (LBR), commonly measured with gradient-echo (GE) MRI sequence, is confounded by non-local changes in deoxygenated hemoglobin and cerebral blood volume propagated within intracortical ascending veins, leading to a unidirectional blurring of the neuronal activity distribution towards the cortical surface. Here, we present a new cortical depth-dependent model of the BOLD response based on the principle of mass conservation, which takes the effect of ascending (and pial) veins on the cortical BOLD responses explicitly into account. It can be used to dynamically model cortical depth profiles of the BOLD signal as a function of various baseline- and activity-related physiological parameters for any spatiotemporal distribution of neuronal changes. We demonstrate that the commonly observed spatial increase of LBR is mainly due to baseline blood volume increase towards the surface. In contrast, an occasionally observed local maximum in the LBR (i.e. the so-called “bump”) is mainly due to spatially inhomogeneous neuronal changes rather than locally higher baseline blood volume. In addition, we show that the GE-BOLD signal laminar point-spread functions, representing the signal leakage towards the surface, depend on several physiological parameters and on the level of neuronal activity. Furthermore, even in the case of simultaneous neuronal changes at each depth, inter-laminar delays of LBR transients are present due to the ascending vein. In summary, the model provides a conceptual framework for the biophysical interpretation of common experimental observations in high-resolution fMRI data. In the future, the model will allow for deconvolution of the spatiotemporal hemodynamic bias of the LBR and provide an estimate of the underlying laminar excitatory and inhibitory neuronal activity.

## Introduction

The fMRI using the blood oxygenation level-dependent (BOLD)^1^ signal provides an indirect, vascular reflection of neuronal activity and, thus, comprises both neuronal and vascular sources of variability (Havlicek et al. (2015), and references therein). Specifically, neuronal activation causes a series of physiological events altering blood oxygenation, including changes in cerebral blood flow (CBF), cerebral blood volume (CBV) and cerebral metabolic rate of oxygen (CMRO2) (Kim and Ogawa, 2012). The resulting decrease in the paramagnetic deoxyhemoglobin (dHb) content leads to reduced magnetic field inhomogeneities within blood vessels and their surroundings and, consequentially, to an increase in MRI signal sensitive to T_2_- or 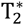-contrasts using spin-echo (SE) or gradient-echo (GE) sequences (Ogawa et al., 1990), respectively. Further, the spatially and temporally varying relationship between these basic physiological processes results in voxel-specific BOLD hemodynamic response functions (Handwerker et al., 2012; Uludag, 2008). Thus, to deduce neuronal processing from measured BOLD signal requires modeling of the relationship between neuronal and hemodynamic processes during both steady-state and transient phases (Buxton et al., 2004; Friston et al., 2003; Havlicek et al., 2015).

FMRI is typically utilized to map brain activity at the macroscopic level with spatial dimensions between 2 and 4 mm. Recent advances in MRI technology and availability of ultra-high magnetic field human scanners (7 T and above) permitted the increasing number of high-resolution fMRI studies at submillimeter voxel (i.e. mesoscopic) resolution (see review on MRI acquisition by Poser and Setsompop (2018)). As a consequence, it is now possible to measure fMRI activation as a function of cortical depth^2^ and potentially study activity changes in histologically-defined cortical layers (De Martino et al., 2013; Fracasso et al., 2018; Kashyap et al., 2017; Kok et al., 2016; Koopmans et al., 2010; Marquardt et al., 2018; Muckli et al., 2015; Olman et al., 2012; Polimeni et al., 2010; Siero et al., 2011). The main motivation for these studies is to investigate the cortical microcircuit during cognitive processes (Douglas and Martin, 2004): Electrophysiological studies showed that feedforward- and feedback-related neuronal computations engage different cortical layers and exhibit clear differences in laminar distribution of neuronal activity (see review by Self et al. (2018) and references therein). Therefore, laminar fMRI promises to non-invasively study the mesoscopic organization of human brain function (see reviews (Dumoulin et al., 2018; Lawrence et al., 2018)). However, data acquisition, analysis and modeling challenges remain to conduct robust neuroscientific studies using laminar fMRI (De Martino et al., 2018; Kemper et al., 2018; Polimeni et al., 2018).

The acquisition method-of-choice often is GE-based BOLD contrast due to its ease of implementation, highest signal-to-noise ratio (SNR) and large brain coverage, but it suffers from spatial confounds related to intra-cortical ascending and pial veins. The GE-based BOLD contrast measures signal mainly from venous vasculature that includes both microvasculature (MV) and macrovasculature (Gagnon et al., 2015; Kim and Ogawa, 2012; Uludağ et al., 2009). As a part of macrovasculature, intracortical ascending veins (AVs) collect the deoxygenated blood from local MV and carry it towards the surface, where it enters larger pial veins (PVs). The anatomical distribution of the vasculature but also related dynamics of blood through different vascular compartments introduce bias in amplitude and localization of observed laminar BOLD response (LBR^3^) with respect to the underlying neuronal activation (see reviews by (Petridou and Siero, 2018; Uludag and Blinder, 2018) and references therein). Thus, AVs introduce cross-talk (aka leakage) between GE-BOLD signals from the different cortical depths.

Until now, there have only been a few attempts to model the effect of AVs on laminar fMRI BOLD signals. Markuerkiaga et al. (2016) proposed a steady-state model using realistic vasculature measured in V1 region in the monkey brain (Weber et al., 2008) in combination with a vascular network model published by Boas et al. (2008) and the fMRI signal model by Uludağ et al. (2009) to predict laminar profiles and point spread functions (PSF) for GE and SE MRI sequences. Their simulation results confirmed that, unlike GE-BOLD contrast, SE-BOLD signal in upper cortical depths has low contamination from lower depths. Furthermore, the simulated LBR of GE-BOLD contrast showed the typical increase towards the surface, reflecting vascular density distribution in both MV and AVs. Within the dynamic causal modeling (DCM) framework (Friston et al., 2003), Heinzle et al. (2016) proposed a two cortical depth model by extending the standard balloon model (Buxton et al., 1998; Friston et al., 2000), linking both depths via AV. In contrast to the model by Markuerkiaga et al. (2016), it is a dynamic model. However, it does not directly model the distribution of vascular density in both MV and AVs, rather it aims to simulate the blood draining from the lower to the upper cortical depth via AV on a phenomenological level. In contrast to the standard balloon model, this two-depth extension does not necessarily comply with the principles of mass conservation for all parameter combinations. Nevertheless, by accounting for strength and delay of the draining effect, the simulated BOLD signal in the two cortical depths compared well (i.e. matching the amplitude and delay of the response peak) with LBRs reported e.g. by Siero et al. (2011).

In the current study, we expanded the balloon model for GE-BOLD signal, such that the total venous signal of each cortical depth is described by a local (venous signal of the local MV) and non-local component (AV carrying dHb concentration changes from the lower depths to the surface). Specifically, we form a multi-compartment cortical depth-dependent vascular model that is (similarly as the original balloon model) defined in terms of absolute and baseline variables and is derived using principles of mass conservation. This also ensures that the proposed model naturally scales for arbitrary number of cortical depths. Additionally, by virtue of being a fully dynamic model described by differential equations, the proposed model not only allows simulating steady-states but also entire time-courses of the LBR, including its dynamic features, such as initial dip, response peak, early-overshoot, or post-stimulus undershoot (PSU).

Our primary aim is to explore the main effects of varying physiological parameters on the resulting LBR and compare these results to experimental observations. Ultimately, beyond testing the physiological hypothesis underlying common observations, we foresee that this new physiological model will allow dynamic deconvolution of the spatial hemodynamic bias of the LBR and provide an estimate of the underlying depth-dependent excitatory and inhibitory neuronal activity (Havlicek et al., 2017b; Havlicek et al., 2019).

**Table 1.** List of frequently used abbreviations.

|  |  |
| --- | --- |
| **BOLD** | **Blood oxygenation level-dependent** |
| **GE** | Gradient echo |
| **CBF** | Cerebral blood flow |
| **CMRO<sub>2</sub>** | Cerebral metabolic rate of oxygen |
| **CBV** | Cerebral blood volume |
| **dHb** | Deoxy-hemoglobin |
| **LBR** | Laminar BOLD response |
| **MV** | Microvasculature |
| **PA** | Pial artery |
| **DA** | Diving artery |
| **AV** | Ascending vein |
| **PV** | Pial vein |
| **PSU** | Post-stimulus undershoot |
| **TTP** | Time to peak |
| **TTU** | Time to undershoot |
| **PTT** | Peak to tail |
| **CBV<sub>0</sub>** | Baseline cerebral blood volume |
| **CBF<sub>0</sub>** | Baseline cerebral blood flow |
| **PSF** | Point spread function |

## Methods

A network of pial arteries (PAs) and PVs covers the surface of the cortex. Diving arteries (DAs) branch off the PAs and penetrate brain tissue supplying oxygenated blood to the cortical parenchyma. Similarly, AVs emerge from the parenchyma and drain partially deoxygenated blood into larger PVs (see illustration in Figure 1A). DAs and AVs (i.e. macrovasculature) are oriented perpendicular to the surface and are connected through a dense network of microvasculature (MV), which has a quasi-random orientation (Uludag and Blinder, 2018). The density of DAs and AVs increases towards the surface; i.e. the baseline blood volume fraction also increases (Duvernoy et al., 1981; Schmid et al., 2017a). The network of MV is formed by arterioles, capillaries (where large majority of oxygen extraction takes place) and venules^4^. Distribution of MV across cortical depth is more homogeneous compared to the macrovasculature (Weber et al., 2008).

**Figure 1.**
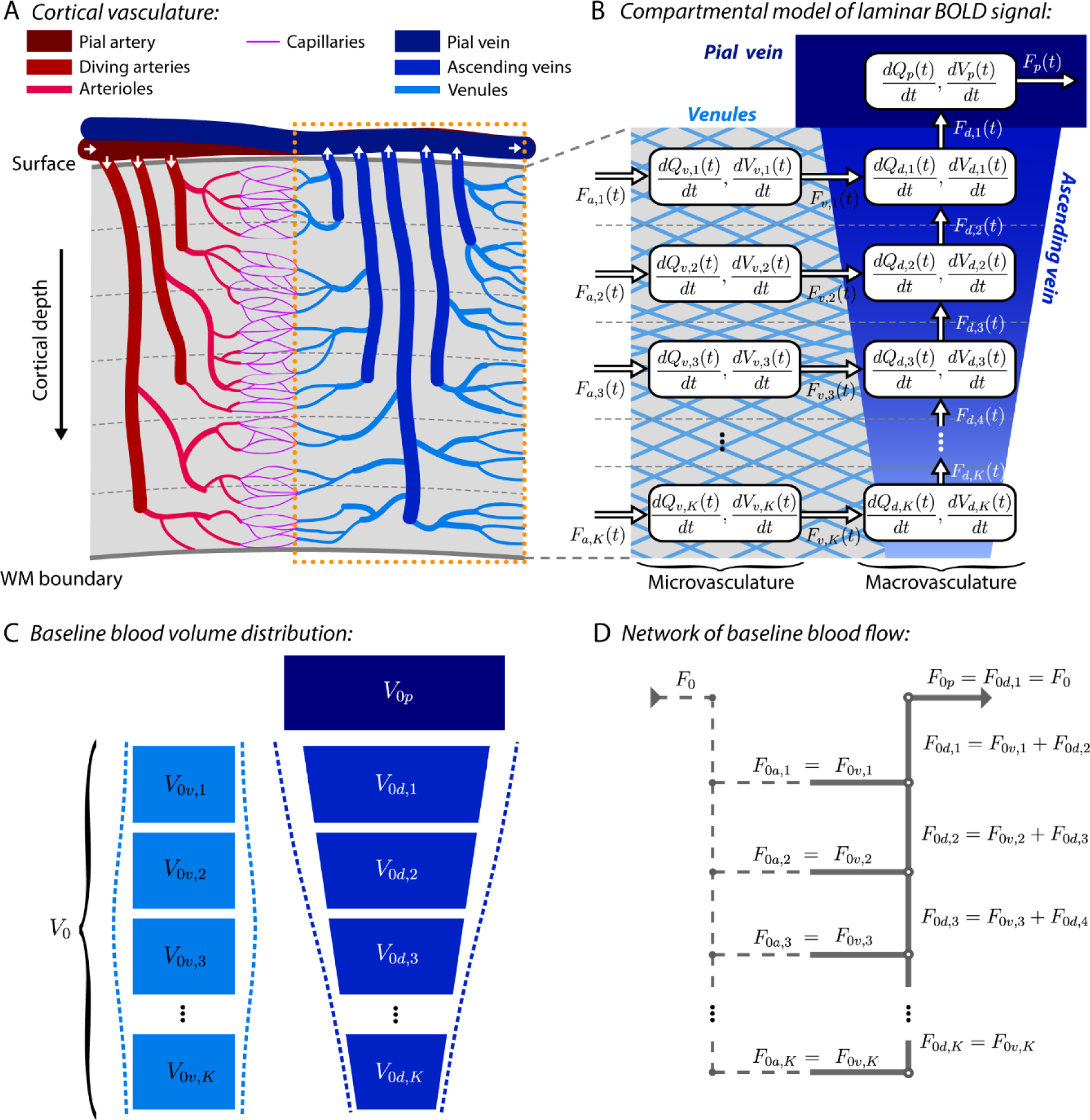
(A) Schematic illustration of depth-dependent distribution of vasculature – from arteries to veins – in the cortex. White arrows indicate direction of the blood flow. Dotted rectangle emphasizes the venous vasculature that mainly contributes to the GE-fMRI signal. (B) Flow-diagram representing the compartmentalized version of the cortical vascular (venous) network, distinguishing microvasculature (venules) and macrovasculature (ascending and pial veins). Compartments are connected with blood flow (*F*) and each compartment models the changes in CBV (*V*) and dHb content (*Q*). (C) Each compartment is further described by baseline CBV. Dashed lines illustrate the possibility to specify diverse laminar distributions of baseline CBV. (D) Network of baseline CBF, representing the merging between venules and ascending vein (based on mass conservation law).

The majority of the BOLD signal acquired with 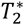-weighted sequences originates in and around post-capillary venous vessels (see Discussion section for details) (Kim and Ogawa, 2012; Uludağ et al., 2009). Within venous vessels, no further oxygen extraction occurs (Boas et al., 2008; Gagnon et al., 2015; Vazquez et al., 2010; Vovenko, 1999). The BOLD signal from a specific cortical depth is affected by local hemodynamic changes in the MV and additionally by the blood carry-over effect towards the surface by the AVs. Note that below we use the terms ‘venules’ and ‘microvasculature’ (MV) interchangeably. The venous vessels and their anatomical organization can be schematized by a simplified network of connected compartments distributed across an arbitrary number of cortical depths. In Figure 1B, vascular compartments are represented by boxes, and the connections between the compartments are depicted by arrows. Every compartment has a number of connections leading to the box (representing blood inflows) and a number of connections leading from the box (representing blood outflows). Within each cortical depth, the GE-MRI-relevant vasculature entails MV and AV. While there are no direct connections between MV compartments across cortical depths, the vertical series of compartments representing the AV is unidirectionally connected with blood flowing towards the cortical surface. The laminar fMRI signal model described below calculates the time courses of dHb and CBV in the MV and AV, respectively. Optionally, one can also consider an additional vascular compartment representing the PV, which is connected to the output of the AV of the top cortical depth. Equations for the PV compartment are included in the Appendix.

### Dynamic changes in hemodynamic variables

Each vascular compartment is described by dynamic changes of blood volume, *dV*_*i*,*k*_(*t*)/*dt*, and dHb content, *dQ*_*i*,*k*_(*t*)/*dt*, where subscript *i* refers to venules or AV (below indicated with subscripts *v* or *d*, respectively) and *k* refers to the *k*-th cortical depth with respect to the pial surface. We use capital letters to represent absolute changes in physiological variables. It is further assumed that the dHb content within each compartment is distributed homogeneously (i.e. the mass is well-mixed) and passed between compartments by following Fick’s principle (i.e. the law of mass conservation). The absolute changes in CBV and dHb of the venules and AV are then governed by the following mass balance equations (for detailed derivation see Supplementary Material 1):

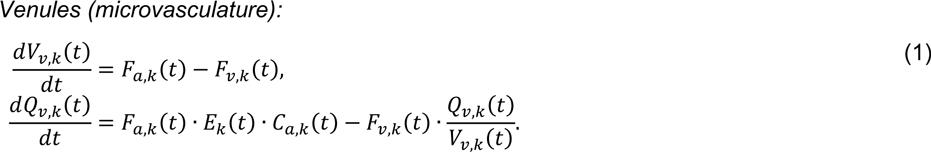

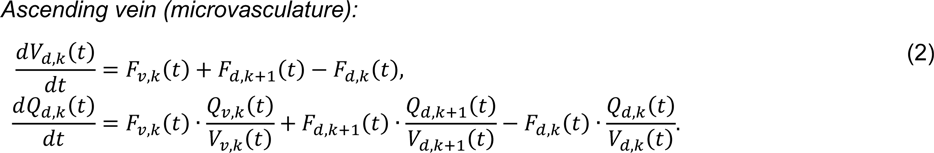

According to the mass conservation law, the difference between the amount of blood flowing in and flowing out is additionally stored in the compartment (i.e. volume change). The equations for the MV are the same as in the original balloon model (Buxton et al., 1998), here only expanded for multiple cortical depths. Therefore, in the venules, the changes in the absolute CBV is a simple difference between the absolute CBF^5^ from the arterioles, *F*_*a*,*k*_(*t*), which is the driving function of the model, and the flow leaving the MV, *F*_*v*,*k*_(*t*). Similarly, the absolute change in the dHb amounts to the difference between the delivery rate of dHb into the venules compartment, *M*_*k*_(*t*) = *F*_*a*,*k*_(*t*) · *E*_*k*_(*t*) · *C*_*a*,*k*_(*t*), i.e. the absolute change in CMRO2 in MV, and the clearance rate of dHb, *F*_*v*,*k*_(*t*) · *Q*_*v*,*k*_(*t*)/*V*_*v*,*k*_(*t*). The change in CMRO2 is the second driving function of the model. Here, *E*_*k*_(*t*) is the net extraction of oxygen from the blood as it passes through the capillary bed, *C*_*a*,*k*_(*t*) is the concentration of oxygen in the arterioles (assuming fully oxygenated blood), and *Q*_*v*,*k*_(*t*)/*V*_*v*,*k*_(*t*) is the dHb concentration in the venules compartments.

For the AV (i.e. Equation 2), the absolute changes in CBV in *k*-th depth are determined by the sum of blood inflows from the venules of the same depth, *F*_*v*,*k*_(*t*), and the lower depth of the AV, *F*_*d*,*k*+1_(*t*), minus the outflow to the upper depth, *F*_*d*,*k*_(*t*) (see also Figure 1B). In the case of the lowest depth *K*, this reduces to the difference between the inflow from venules and outflow to the upper depth of the AV. The absolute changes of dHb in the AV are again described by the mass balance between the delivery rate of dHb into the compartment (i.e. from venules and possibly also from lower depth of AV) and wash out rate of dHb out from the compartment.

Next, it is common to express the above dynamic equations in terms of normalized (relative) variables with respect to their baseline values: *f*_*i*,*k*_(*t*) = *F*_*i*,*k*_(*t*)/*F*_0*i*,*k*_, *m*_*k*_(*t*) = *M*_*k*_(*t*)/*M*_0_, *q*_*i*,*k*_(*t*) = *Q*_*i*,*k*_(*t*)/*Q*_0*i*,*k*_, and *v*_*i*,*k*_(*t*) = *V*_*i*,*k*_(*t*)/*V*_0*i*,*k*_. Hereafter, we use subscript 0 to indicate baseline values. The laminar hemodynamic model formulated in terms of relative variables is presented in the Appendix. Specific assumptions about the laminar distribution of baseline blood volume (CBV_0_) and flow (CBF_0_) are briefly described below and in more detail in the Supplementary Material 1. Further, the steady-state relationship between CBF and CMRO2 in the MV is described by (possibly) depth-specific *n*-ratio, *n*_*k*_ = (*f*_*a*,*k*_ − 1)/(*m*_*k*_ − 1) (Buxton et al., 2004). Please note that CBF can also be dynamically uncoupled with respect to CBV and CMRO2 during transient periods (Chen and Pike, 2009; Frahm et al., 2008; Mandeville et al., 1999; van Zijl et al., 2012).

### CBF-CBV relationship

The relative change in the blood flow leaving the compartments depends on the volume change. Thus, for both venules and AV compartments, we assume a power-law relationship between blood outflow and volume during steady-state and additional dynamic viscoelastic effect (accounting for the CBF-CBV uncoupling) during transient states:

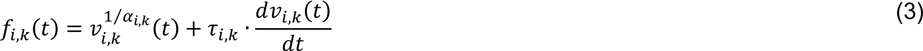

Here *α*_*i*,*k*_ is an exponent relating CBV to CBF response during steady-state (Grubb et al., 1974) and *τ*_*i*,*k*_ is the viscoelastic constant that controls how fast the CBV adjusts to the change in the CBF through the compartment. Both *α*_*i*,*k*_ and *τ*_*i*,*k*_ are defined as potentially being vascular compartment- and depth-specific. In addition, the possibility that the viscoelastic constants differ during inflation and deflation could be also included by choosing different constants during inflation and deflation (e.g., *τ*_*i*+_ and *τ*_*i*−_). Further, in the case of no CBV change in the compartment, the outflows from MV or AV take the following form (as the above relation does not hold for *α*_*i*_ = 0):

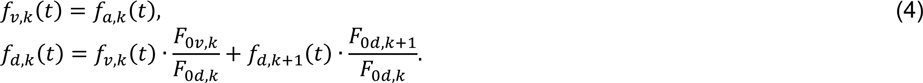

The relative outflow from *k*-depth of AV is given by a weighted sum of inflow from MV in the same depth and inflow from lower depth of the AV. The weighting factors then account for differences in baseline CBF between compartments, which is a result of mass conservation law (see below).

### Baseline blood volume and flow

For a macroscopic model that describes a brain region by a single vascular compartment, one needs to define the total amount of CBV_0_ (i.e. *V*_0_ in equations above) and CBF_0_ (i.e. *F*_0_ in equations above). For a laminar model, we have a network of connected compartments. Thus, we need a cortical-depth-specific and vasculature-type-specific distribution of CBV_0_ (*V*_0*i*,*k*_) and CBF_0_ (*F*_0*i*,*k*_), reflecting our knowledge about the anatomy of the vascular organization, as described above (see Figure 1C and D). Depth-specific distribution of CBV_0_ in the MV and AV can be freely defined, but summing up CBV_0_ from all depths has to yield the total amount of CBV_0_. We express the absolute baseline CBV in units of mL. In simulations described below, we are mostly concerned with linear increase of CBV_0_ towards the surface either in MV or AV. For this reason, we have parameterized the slope of the increase of CBV_0_ towards the surface with a positive constant *s*_*i*_ (detail description is provided in Supplementary Material 1). This ensures that model complexity does not increase with a higher number of cortical depths, and because the slope *s*_*i*_ is independent of number of depths, the model is easily scalable to (almost) arbitrary number of depths. However, in the following text, we also characterize the linear increase of CBV_0_ towards the surface with a ratio between CBV_0_ in the superficial and the lowest depth, *V*_0*i*,1_/*V*_0*i*,*K*_ (note that this value is number of depth-dependent).

The CBF_0_ is distributed between compartments as illustrated in Figure 1D. The number of cortical depths defines the number of junctions, where baseline flow splits from DA to arterioles, and equally the number of junctions, where baseline flow in venules merges within the AV. Splitting and merging of the baseline flow follow the mass conservation law (i.e. continuity equation in particular). That is, the CBF_0_ in *k*-th depth is a sum of blood flow leaving the MV of that depth, and the blood flow from the lower depth (i.e. *k* + 1) of the AV (see Figure 1D). Consequently, the blood flow leaving the AV in the most superficial depth equals the total amount of CBF_0_ in the MV. As the CBF_0_ in AV increases towards the surface, so must also the CBV_0_ to maintain physiologically plausible transit times through AV. We express the absolute baseline CBF in units of mL per sec. At any point of the vascular tree, the mean transit time of blood through a specific vascular compartment (in sec) is defined as the ratio of CBV_0_ to CBF_0_, *t*_0*i*_ = *V*_0*i*,*k*_/*F*_0*i*,*k*_ (called the central volume principle). Note that for relatively large (linear) increase of CBV_0_ towards the surface in the AV (e.g. for six depths if *V*_0*d*,1_/*V*_0*d*,6_ = 6), the transit times through AV remain constant across depths. To introduce some decrease in transit time as we move closer to the surface, which is known from invasive measurements of vascular physiology (Schmid et al., 2017b) (i.e. the blood is drained faster from upper depths due to larger increase in CBF_0_ than CBV_0_), one can consider smaller ratio (e.g. *V*_0*d*,1_/*V*_0*d*,6_ = 3). Additional details on CBF_0_, CBV_0_, transit time and their dependence on the number of depths are provided in Supplementary Materials 1 and 2.

### Laminar BOLD signal equation

The baseline BOLD signal for GE sequence in *k*-th depth is described as a CBV_0_-weighted sum of extra- (*E*) and intra-vascular (*I*) signals originating from the tissue around and blood inside venules and AVs, respectively:

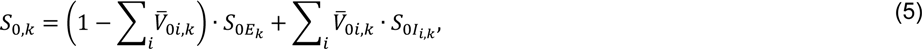

Where 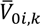 is the CBV_0_ fraction with respect to GM tissue in the *k*-th depth of the venules or AV compartment (again denoted by subscript *i*). The transformation from CBV_0_ in absolute units (i.e. in mL) to fractions is provided in Supplementary Material 1. The baseline extra- and intra-vascular signal components can be written as 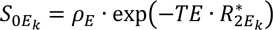 and 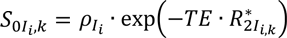, where *TE* is the echo-time, 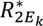 and 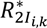 are the corresponding baseline transverse relaxation rates, and *ρ*_*E*_ and *ρ*_*Ii*_ are water proton densities in GM tissue and blood, respectively. *T*_1_ effects are neglected in this model (see also Supplementary Material 1) but can be easily included. Below, we assume that the baseline relaxation rates are constants across depths, but this assumption can also be easily relaxed (see Discussion section).

During neuronal activation, the transverse relaxation rates of the extra- and intra-vascular signals in different compartments are altered by depth-specific increments 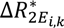 and 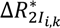, respectively, and the venous (absolute) CBV changes to a new value 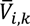. Then, by: (1) using a first-order Taylor expansion to approximate the change in BOLD signal from baseline to activation, Δ*S*_*k*_ (Stephan et al., 2007); (2) relating the changes in transverse relaxation rates of extra- and intra-vascular signals to absolute changes in dHb content, *Q*(*t*), and concentration, *Q*(*t*)/*V*(*t*), respectively (Boxerman et al., 1995; Obata et al., 2004; Ogawa et al., 1993); and (3) expressing the dHb content and CBV in terms of their relative changes with respect to the baseline values, as defined in the compartmental model above, the fractional change of the BOLD signal in *k*-th depth is (detailed derivation is included in the Supplementary Material 1):

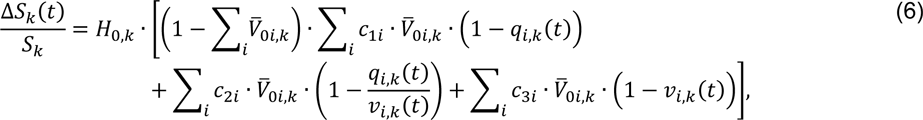

where 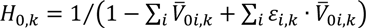. The three constants, *c*_1*i*_, *c*_2*i*_, and *c*_3*i*_ scale the relative changes in extra-, intra-vascular signals and CBV, respectively.

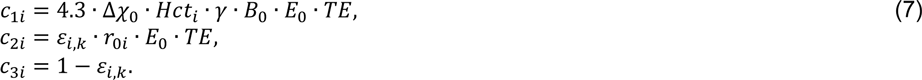

They describe the BOLD signal dependence on physical MR and physiological parameters: magnetic field strength, *B*_0_; echo-time *TE*; susceptibility difference between fully oxygenated and deoxygenated blood, Δ*χ*_0_; gyromagnetic ratio of water protons, *γ*; slope of change in intravascular signal relaxation rate with change in oxygen saturation, *r*_0*i*_; baseline oxygen extraction fraction, *E*_0_; proportion of hematocrit in the blood, *Hct*_*i*_; and ratio of baseline intra-to-extra-vascular signals, *ε*_*i*,*k*_ = *S*_0*I_i,k_*_/*S*_0*E_k_*_. One can notice that some of these parameters can be defined as vascular compartment- and/or cortical depth-specific. Values and plausible ranges of these parameter related to different vascular compartments for 7 T magnetic field and GE sequence are reported in Table 2. Note that this provides an updated list of parameters that we proposed earlier (Havlicek et al., 2015).

**Table 2.**
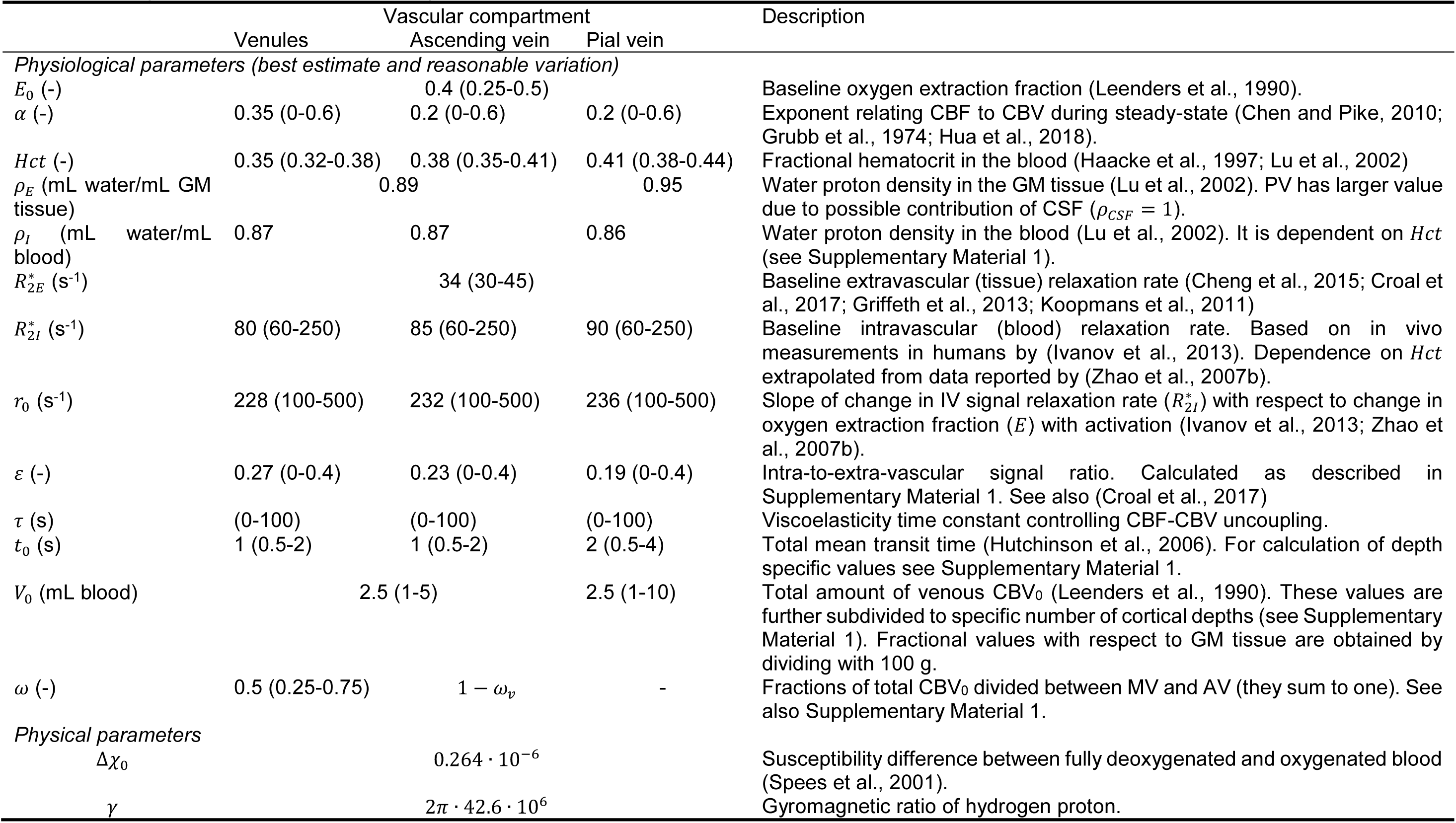
Model parameters for 7 T and GE sequence and TE ≅ 28 ms.

## Simulations

The dynamic nature of our new laminar BOLD model allows us to explore the dependence of the modeled LBR on different physiological parameters during both steady-state and transient periods. Thus, below we choose certain parameter sets, which allow us to characterize the properties of the model or to mimic certain experimental observations. Additionally, we also demonstrate the utility of the new model by exploring physiologically plausible scenarios for a recently published study with depth-specific BOLD signal and CBV changes in the human motor cortex (Huber et al., 2017). However, as the examples below illustrate plausible yet specific scenarios for some common observations in high-resolution fMRI and optical imaging, we encourage readers to further explore the model for other parameter combinations and experimental observations using our MATLAB code available at https://github.com/martinhavlicek/laminar_BOLD_model.

All simulations were performed in MATLAB R2015b (The MathWorks Inc., Navick, USA), and the differential equations describing the laminar BOLD model were evaluated using Euler’s method (with integration step, Δ*t* = 0.01 s). The following paragraphs provide details on how the specific simulations, later shown in the Results section (labeled with corresponding subheadings), were performed.

### Steady-state simulations

The most characteristic observation on LBR dependence acquired with GE sequence is that it gradually increases from lower to upper depths (Uludag and Blinder (2018) and references therein). The LBR increase is expected to depend on brain region, stimulus-type and duration, voxel-selection, but also on MR parameters and sequence. Therefore, there is a wide range of reported values on BOLD signal amplitude variation between lower depths (can be as low as ∼0.5%) and upper depths (can be as high as ∼12%) (Kashyap et al., 2017; Polimeni et al., 2010; Siero et al., 2011). Some studies report simple or superlinear increase of the amplitude towards the surface (De Martino et al., 2013; Kashyap et al., 2017), while others also show some non-monotonic behavior (generally described as ‘bumps’) in the LBR (Chen et al., 2013; Huber et al., 2017; Koopmans et al., 2010; Marquardt et al., 2018).

First, we define a default scenario of physiological assumptions inspired by experimental data and, by variation of some of the assumptions, we then explore the main factors affecting the increase of LBR towards the surface. Second, we describe simulation scenarios that are motivated by specific experimental observations, commonly made in laminar studies, for which, however, different physiological explanations have been put forward.

#### Default scenario

We divide the cortex into six cortical (equivolume) depths (*K* = 6), where the first and sixth indices refer to the upper (i.e. by the cerebrospinal fluid (CSF) boundary) and lower (by the white matter (WM) boundary) depths, respectively. The total amount of venous CBV_0_ is 2.5 mL, from which 50% relates to MV (*ω*_*v*_ = 0.5) and 50% to AV (*ω*_*d*_ = 0.5) (Weber et al., 2008). Next, we choose that CBV_0_ distribution is constant across depths in the MV but increases towards the surface in the AV with a ratio between CBV_0_ in the superficial and the lowest depth of AV, *V*_0*d*,1_/*V*_0*d*,6_ = 3 (i.e. *s*_*d*_ = 0.3). The transit time through MV is kept constant, *t*_*v*0_ = 1 s (Hutchinson et al., 2006), and the transit times through AV are calculated automatically, as described above using the central volume principle (i.e. in this case, decreasing towards the surface). In terms of functional changes (during activation), we assume that in each depth there is a 60% increase of relative CBF in the MV accompanied with changes in CMRO2 that are linearly coupled to CBF (with *n* = 4). Further, we assume CBV changes in the MV and AV of ∼18% and ∼10%, defined by *α*_*v*_ = 0.35 and *α*_*d*_ = 0.2, respectively (Hua et al., 2018; Stefanovic et al., 2008). Finally, we assume that 65% of oxygen is extracted upon the arrival to venules (*E*_0_ = 0.35), with no further oxygen extraction along venous vessels (i.e. assuming oxygen saturation of hemoglobin (1 − *E*_0_) being homogeneously distributed across cortical depths). The laminar BOLD signal equation is parameterized for GE sequence, with *B*_0_ = 7 T and *TE* = 28 ms.

#### Main factors affecting LBR increase towards surface

Starting from the default scenario, we examined the following cases: (i) the effect of increase of CBV_0_ distribution towards the surface (*V*_0*i*,1_/*V*_0*i*,6_ = 3, i.e. with slope *s*_*i*_ = 0.3) vs constant distribution (*V*_0*i*,1_/*V*_0*i*,6_ = 1, i.e. with slope *s*_*i*_ = 0) in either MV or AV; (ii) changing CBV_0_ ratio between MV and AV (*V*_0*v*_/*V*_0*d*_ = 1, 0.5 or 2); (iii) the effect of increasing relative CBF change (20%; 60% and 100%). Other effects, such as increasing relative CBV change in the MV or AV vs no CBV change or changing the *n*-ratio between relative CBF and CMRO2 were explored as well (see Supplementary Material 2). All these scenarios were simulated by assuming uniform neuronal activation (i.e. same CBF increase) across all depths.

Additionally, for selected scenarios, we derive laminar point spread functions (PSFs) by activating individually each cortical depth (Markuerkiaga et al., 2016). The laminar PSF is generally described by a peak signal increase in the activated depth, followed by a lower amplitude tail representing a signal leakage towards the upper depths. Thus, to explore how different physiological scenarios affect the relative leakage of the signal, we also calculated peak-to-tail (PTT) ratio. Following Markuerkiaga et al. (2016), the PTT is defined as the ratio between the PSF peak amplitude and an average tail amplitude over all depths towards the surface.

#### Neuronal vs vascular origin of ‘bump’ in LBR

In some studies, a local maximum of the BOLD signal amplitude in the middle depths has been observed, which has either been attributed to be of neuronal or vascular origin (Chen et al., 2013; Goense and Logothetis, 2006; Harel et al., 2006; Koopmans et al., 2010). The neuronal hypothesis postulates that the observed relative maximum in the LBR reflects variation in the neuronal signal, which should then also be directly reflected in the laminar profile of absolute CBF. In contrast, the vascular hypothesis proposes that the observed variation in the LBR is due to non-uniform distribution of baseline vascular density across depths (Lauwers et al., 2008). The current model allows us to test these two (non-exclusive) hypotheses.

We simulated the *neuronal hypothesis* by considering variable absolute CBF change with a clear bump in the middle (40% larger) compared to its adjacent depths, assuming that higher neuronal activity is related to larger increase in CBF (see Figure 2A). The CBF_0_ and CBV_0_ in MV were considered constant across cortical depths. The *vascular hypothesis* was simulated by assuming variable CBV_0_ that was 40% larger in the middle compared to its adjacent depths (see Figure 2B). This is a representative range (albeit at the higher end) of possible variation of CBV_0_ in the MV (for example, in macaque V1 (Weber et al., 2008)), and it is comparable to variation (i.e. also 40%) in absolute and relative CBF used in the neuronal hypothesis. Further, in this case, the absolute CBF change was constant across depths and the relative CBF change was then calculated as defined above. Moreover, we have also evaluated a *mixed hypothesis*, as both neuronal and vascular effects can be present in actual experiments. Instead of considering the absolute CBF to be constant, we introduced variation in the absolute CBF change (i.e. neuronal variation) in a way that it matches the size of variation in the CBF_0_, so that the relative CBF change is constant for all depths (see Figure 2C). Thus, in this case, there are colocalized relative maxima at both neuronal and vascular levels.

**Figure 2.**
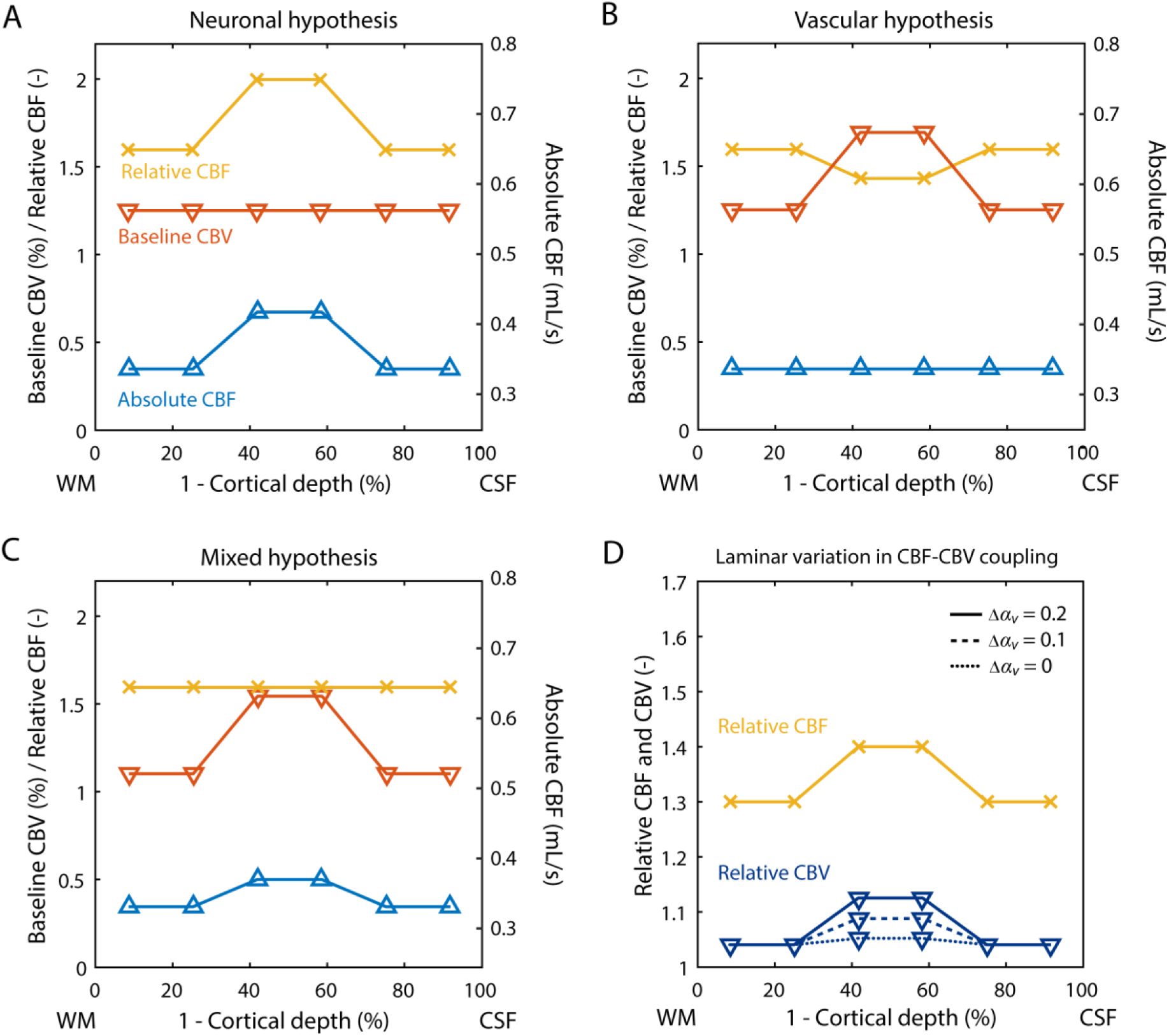
Simulation scenarios demonstrating variation of physiological variables across depths. Each plot depicts physiological variables as a function of (1 – normalized cortical depth) between WM and CSF boundaries. (A-C) Comparison of three simulated physiological hypotheses underlying the observed bump in the LBR. Distinct laminar profiles of CBV_0_, relative and absolute CBF in the MV are displayed for: (A) neuronal, (B) vascular, and (C) mixed hypotheses. (D) Simulations demonstrating variable CBF-CBV coupling across cortical depths in the MV. For relative CBV, three different coupling parameters (*α*_*v*_) with respect to CBF in the middle depths are shown, differing from surrounding depths by Δ*α*_*v*_.

#### Laminar variation in CBF-CBV relationship

Some animal studies observed that the steady-state CBF-CBV relationship, generally characterized by the exponent, *α*, may vary across cortical depths (Jin and Kim, 2008; Zhao et al., 2007a). Note that these observations also included variable CBF change across depths as well. Therefore, to explore this scenario and its impact on the LBR, we have considered variable relative CBF change across depths (similarly as in the neuronal hypothesis but assuming smaller between depths variation of 10% to more closely match the above-mentioned experimental observations) and additionally varied the strength of CBF-CBV relationship (i.e. *α*_*v*_) in different depths of MV (see Figure 2D). In the lower and upper depths, we fixed *α*_*v*_ = 0.15, and in the middle depths, we varied an increase in CBF-CBV coupling, Δ*α*_*v*_, between 0 and 0.2 (i.e. referring to a larger CBV change in the middle depths compared to surrounding depths). Possible scenarios of variable CBF-CBV coupling across depths in AV or in both MV and AV were explored as well.

### Simulations of response transients

It is typically observed that the amplitude of response transients (relative to the positive peak), such as the initial dip, the early-stimulus overshoot, response adaptation and PSU, increases towards the upper depths (see (Uludag and Blinder, 2018), and references therein). Similarly, the delay of BOLD response transients that can be described by time-to-peak (TTP) and time-to-undershoot (TTU) most often increases towards the superficial depths as well (see (Petridou and Siero, 2018), and references therein). In general, BOLD response transients can be evoked by different dynamic relationships between physiological mechanisms. First, they can represent close reflection of neuronal dynamics (i.e. having neuronal origin), likely then observed also in the CBF response (Bandettini et al., 1997; Havlicek et al., 2015; Mullinger et al., 2013; Sadaghiani et al., 2009; Shmuel et al., 2006). Second, they can result from dynamic uncoupling between CBF and CBV that can appear either in the micro- or macro-vasculature (i.e. having vascular origin) (Buxton et al., 1998; Chen and Pike, 2009; Havlicek et al., 2017a; Mandeville et al., 1999). Third, they can theoretically also be due to dynamic uncoupling between CBF and CMRO2 in the MV (i.e. having metabolic origin) (Donahue et al., 2009; Frahm et al., 2008; Lu et al., 2004; van Zijl et al., 2012).

The new laminar BOLD model can simulate any of the physiological mechanisms underlying response transients: The neuronal and metabolic origins of response transients can be defined directly by specifying model inputs in terms of CBF and CMRO2 responses. The vascular origins (i.e. the CBF-CBV uncoupling) are controlled independently for the MV and AV via viscoelastic time constants *τ*_*v*_ and *τ*_*d*_, respectively. Unless specified otherwise, we assume the same default parameter setting of the laminar BOLD model as described above for the default scenario of the steady-state.

#### Coupled vs uncoupled dynamic relationships

First, we consider two illustrations of LBR to short 2 s stimuli in order to provide a basic idea of how response transients can be affected by specific physiological assumptions in the laminar model. In the first example, we assume that both CMRO2 and CBV responses are tightly coupled with CBF responses. The stimulus induces the same positive CBF response in all depths, which increases sharply, shortly after the stimulus onset, and starts to gradually decrease to the baseline shortly after the stimulus offset. The second example considers uncoupling and more complex time courses of the basic physiological variables in order to elaborate on possible physiological scenarios of experimental observations. That is, we assume that CMRO2 response increases before the CBF response during the early response phase, then they synchronize (assuming default *n*-ratio) and later they both exhibit a small PSU. Additionally, we consider a small CBF-CBV uncoupling during both inflation and deflation phase in the MV (*τ*_*v*+_ = *τ*_*v*−_ = 2 s). This is the same in the AV, during the inflation phase but larger CBF-CBV uncoupling is assumed during the deflation phase (*τ*_*d*+_ = 2 s and *τ*_*d*−_ = 20 s). Note that *τ*_*d*−_ (or *τ*_*v*−_) may be proportional to the duration of full recovery of PSU of BOLD response, which can last even for very short stimuli between 10 to 30 s (Hirano et al., 2011; Hu et al., 1997).

#### Main factors affecting LBR transients

By first considering the neuronal origin of BOLD response transients, we examined their cortical depth-dependence with respect to: (i) the effect of increased CBV_0_ towards the surface in the MV or AV; (ii) different CBV_0_ ratio between MV and AV; and (iii) the effect of changing transit time through MV. In contrast to the two scenarios above, we assumed input CBF response to 20 s stimulus duration, which additionally included early-overshoot (reaching 60% signal change, as in the case of short stimulus or steady-state scenario) followed by slow adaptation to steady-state plateau (reaching 50% signal change) and small PSU (−10% signal change). Note, that the neuronal and metabolic origins of BOLD response transients showed very similar effects in terms of cortical depth-dependence, and therefore, are not considered here separately.

Next, we examined the dependence of LBR transients being purely of vascular origin. That is, we evaluated the effect of vascular uncoupling between CBF and CBV in the MV and/or AV (*τ*_*v*_ = 20 and/or *τ*_*d*_ = 20 s, same both inflation and deflation phase). The input CBF response was assumed to be the same across depths (i.e. reaching ∼60% signal change) but without early-overshoot and PSU. The CBF-CBV coupling during steady-state was chosen to be identical for both MV and AV (i.e. *α*_*v*_ = *α*_*d*_ = 0.3). All the other parameters in these simulations followed the default scenario. For all these simulations, we have evaluated cortical depth-dependence of early-stimulus overshoot and PSU by examining their amplitudes, TTP and TTU. The TTP and TTU were calculated with respect to stimulus onset and offset, respectively (Jin and Kim, 2008).

### Simulations illustrating laminar fMRI data by Huber et al. (2017)

In the following, we take inspiration from a high-resolution laminar study done by Huber et al. (2017) to demonstrate the ability of the laminar model to simulate, describe and explain complex experimental data. Huber et al. simultaneously measured activity changes in GE-BOLD signal and total CBV using a slice-saturation slab-inversion vascular-space-occupancy (SS-SI-VASO) sequence (Huber et al., 2014) in the primary motor cortex during sequential finger tapping (see Fig.2F in (Huber et al., 2017)). Their laminar profiles were sampled from 21 equivolume depths. The average laminar profile of total CBV during positive response clearly showed two peaks in the lower and upper depths, which had approximately the same amplitude. Note that the total CBV reflects contributions from both arterial and venous CBV, originating mainly in the MV and was reported quantitatively in units of mL. In contrast, LBR showed typical amplitude increase within the grey matter from ∼2% at the WM boundary to ∼10% at the CSF boundary, with the peaks in the CBV profile not or only weakly reflected in the BOLD signal amplitudes.

We examined whether their results (i.e. the laminar discrepancy between total CBV and the BOLD signal) are consistent with our model predictions. To this end, we simulated laminar responses in 21 depths, assuming that the steady-state relative CBF response (i.e. the model input) during 30 s stimulation period has a similar laminar profile as the total CBV reported by Huber et al. (2017)^6^. To simulate a range of LBR between lower and upper depths that is closely comparable with the experimental data, we manipulated the amplitude of relative CBF response, the slope of CBV_0_ in AV, and CBV changes in both MV and AV. After systematically exploring the parameter space, we chose a physiologically plausible scenario (by taking into account also plausible behavior of dynamic response features – see below): relative CBF reaching ∼60% in both lower and upper peaks and ∼45% between them; increasing CBV_0_ in the AV towards the surface (*V*_0*d*,1_/*V*_0*d*,21_ ≅ 4.5, i.e. with slope *s*_*d*_ = 0.4); and CBV changes in the MV (*α*_*v*_ = 0.25) and AV (*α*_*d*_ = 0.1). The other parameters were kept the same as described for the default scenario of the steady-state response.

Next, to further illustrate cortical depth-dependence of dynamic response features in this scenario with 21 cortical depths, we additionally introduced a small response adaptation during stimulation and small PSU in the CBF response. The two-peak laminar activity profile of the CBF response was reflected also during these transient periods. In terms of dynamic vascular features, we have assumed smaller CBF-CBV uncoupling in the MV (*τ*_*v*+_ = *τ*_*v*−_ = 10 s) and larger CBF-CBV uncoupling in the AV (*τ*_*d*+_ = 40 s; *τ*_*d*−_ = 100 s), specified separately for inflation and deflation phases.

## Results

The following paragraphs describe the results of simulations that were in detail specified above and are labeled with corresponding subheadings.

### Results for steady-state simulations

#### Main factors affecting LBR increase towards surface

Figure 3A shows the LBR as a function of CBV_0_. For homogenous CBV_0_ in both MV and AV, the LBR is flat (dotted line). For linear increase of CBV_0_ in the MV or the AV towards the surface, the LBR exhibits a linear increase as well (indicated by dashed and solid lines, respectively). The same spatial increase of CBV_0_ in the MV as in the AV (with a total amount of CBV_0_ being equally divided between MA and AV) results in a smaller LBR increase towards the surface. Next, Figure 3B shows the effect of different CBV_0_ ratios between MV and AV (with default CBV_0_ increase, *V*_0*d*,1_/*V*_0*d*,6_ = 3). Larger CBV_0_ occupied by AV (dashed line in Figure 3B) increases the relative slope of LBR towards the surface.

**Figure 3.**
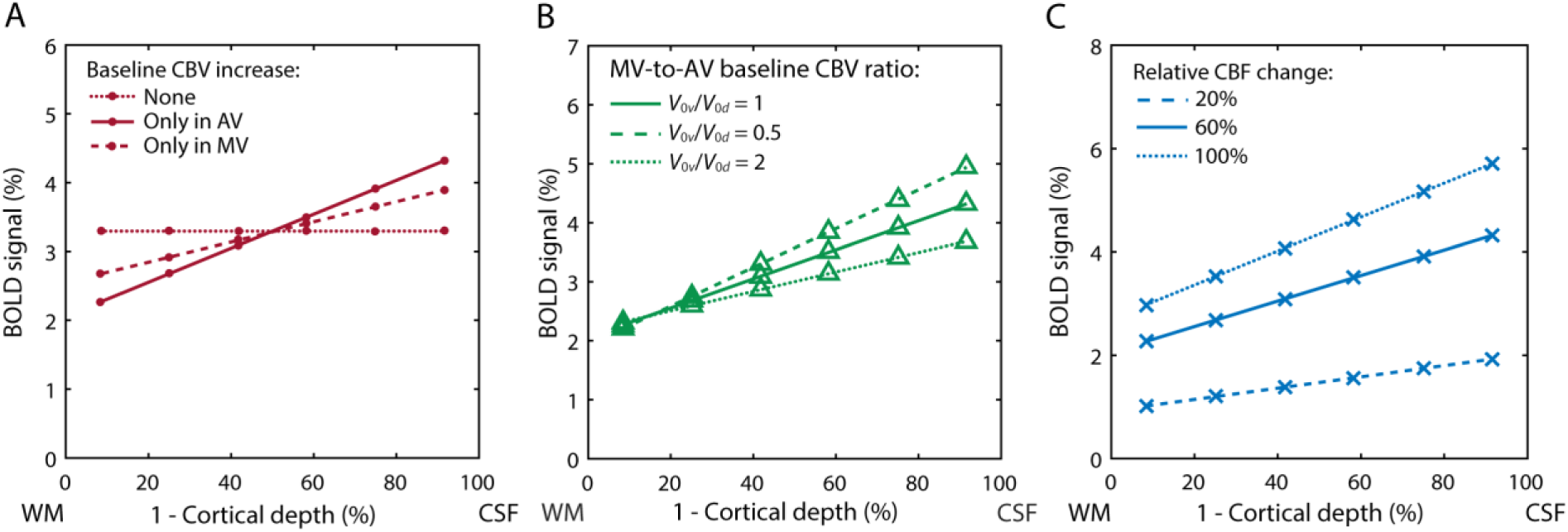
Simulated LBRs during steady-state depicted as function of (1 – normalized cortical depth) between WM and CSF boundaries. (A) LBR dependence on CBV_0_ increase towards the surface (*V*_0*i*,1_/*V*_0*i*,6_ = 3) in different vascular compartments. Constant CBV_0_ across cortical depths in both MV and AV results in a flat LBR. (B) LBR dependence on CBV_0_ ratio between MV and AV, assuming default spatial CBV_0_ increase in the AV. (C) LBR dependence on different amplitudes of relative CBF change in the MV (equal at each depth) – with increasing relative CBF, the amplitudes of the LBR are scaled equally for all depths.

The impact of laminar CBV_0_ distribution can be further modulated by other physiological factors, i.e. changes in CBF, CBV, or CMRO2. Figure 3C shows that with increasing amplitude of the CBF response the amplitudes of LBR increase. Specifically, signal amplitudes in both lower and upper depths scale almost equally; i.e. the relative slope of LBR is almost the same. We refer to this as a simple scaling effect. Similar simple scaling effects on LBR are also obtained by changing the amplitudes of CBV change in the AV or CMRO2 change in the MV (see Figure S2.1 in Supplementary Material 2).

In Figure 4A, we show how the slope of CBV_0_ increase in the AV affects laminar PSFs: (i) with default CBV_0_ increase towards the surface (*V*_0*d*,1_/*V*_0*d*,6_ = 3, depicted with solid lines), the maxima of individual PSFs and amplitude of PSF tails slightly decreases towards the surface; (ii) with larger CBV_0_ increase (*V*_0*d*,1_/*V*_0*d*,6_ = 6, depicted with dashed lines), the maxima and amplitude of tails are slightly increasing towards the surface. Note that, in the first case, the transit time through individual compartments of the AV decreases towards the surface, whereas in the second case, the transit times are constant. Additionally, for the two CBV_0_ distributions, dependence of the average PTT ratio on relative CBF change is depicted in Figure 4B. As expected, with a larger increase of CBV_0_ towards the surface, the PTT ratio is smaller, which means that there is more relative signal leakage towards upper depths (dash line). Importantly, in both cases, the PTT is dependent on the amplitude of the relative CBF change. Specifically, PTT ratio drops (sublinearly) by ∼25% between 20 and 80% of CBF change. This means that with increased neuronal activity there is more relative signal leakage from lower to upper depths. Within the Supplementary Material 2, we also show that a constant CBV_0_ in the AV (*V*_0*d*,1_/*V*_0*d*,6_ = 1) together with a variable laminar CBF response across depths represented by two peaks in the lower and upper depths, may result in lower amplitude of LBR in the upper depths compared to the lower depths (see Figure S2.3). This is due to much shorter transit time in upper depths compared to the lower depths with constant CBV_0_ in the AV. That is, any modification of physiological and anatomical assumptions, including the strength of depth-specific neuronal activation, results in changes in laminar PSFs.

**Figure 4.**
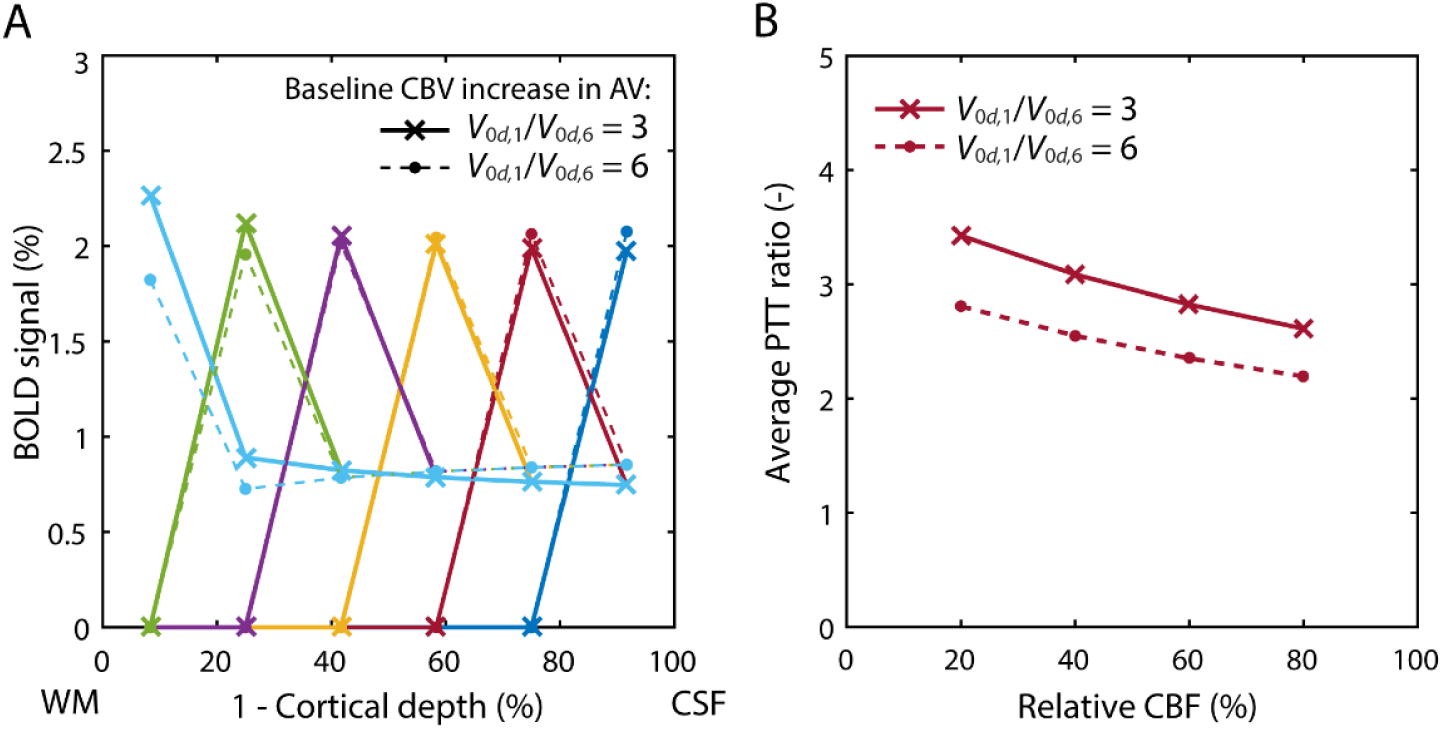
(A) Laminar PSFs for two different slopes of CBV_0_ increase towards the surface in the AV, corresponding to *V*_0*d*,1_/*V*_0*d*,6_ = 3 and *V*_0*d*,1_/*V*_0*d*,6_ = 6 (parameterized with *s*_*d*_ = 0.3 and *s*_*d*_ = 1.2), respectively. The PSF peak appears always in the activated depth, followed by the tail (describing the amount of signal leakage) towards upper depths. (B) For the same CBV_0_ distributions, dependence of the average PTT ratio on relative CBF change.

#### Neuronal vs vascular origin of bump in LBR

In Figure 5A, we show that the neuronal hypothesis yields a considerable BOLD signal bump in the middle depths on top of the typical increase towards the surface (solid line). One can also notice that higher neuronal/CBF increase in the middle depths results in a significant increase of LBR amplitudes in the upper depths; i.e. diverging from the linear trend (depicted with a thin line) predicted for LBR with homogenous distribution of CBF response across depths (as shown above in Figure 3). In contrast, the vascular hypothesis (dotted line) yields almost negligible deviation from the linear increase of the LBR in the middle depths. This is because the amplitude of the BOLD signal is affected by two competing effects: scaling of the BOLD signal amplitude by higher CBV_0_ in the middle depths accompanied by a reduced relative CBF change. The mixed hypothesis (dashed line) also yields a bump in the LBR. Additional factors that could further contribute to the observed bump in the middle cortical depths are described in the Discussion section.

**Figure 5.**
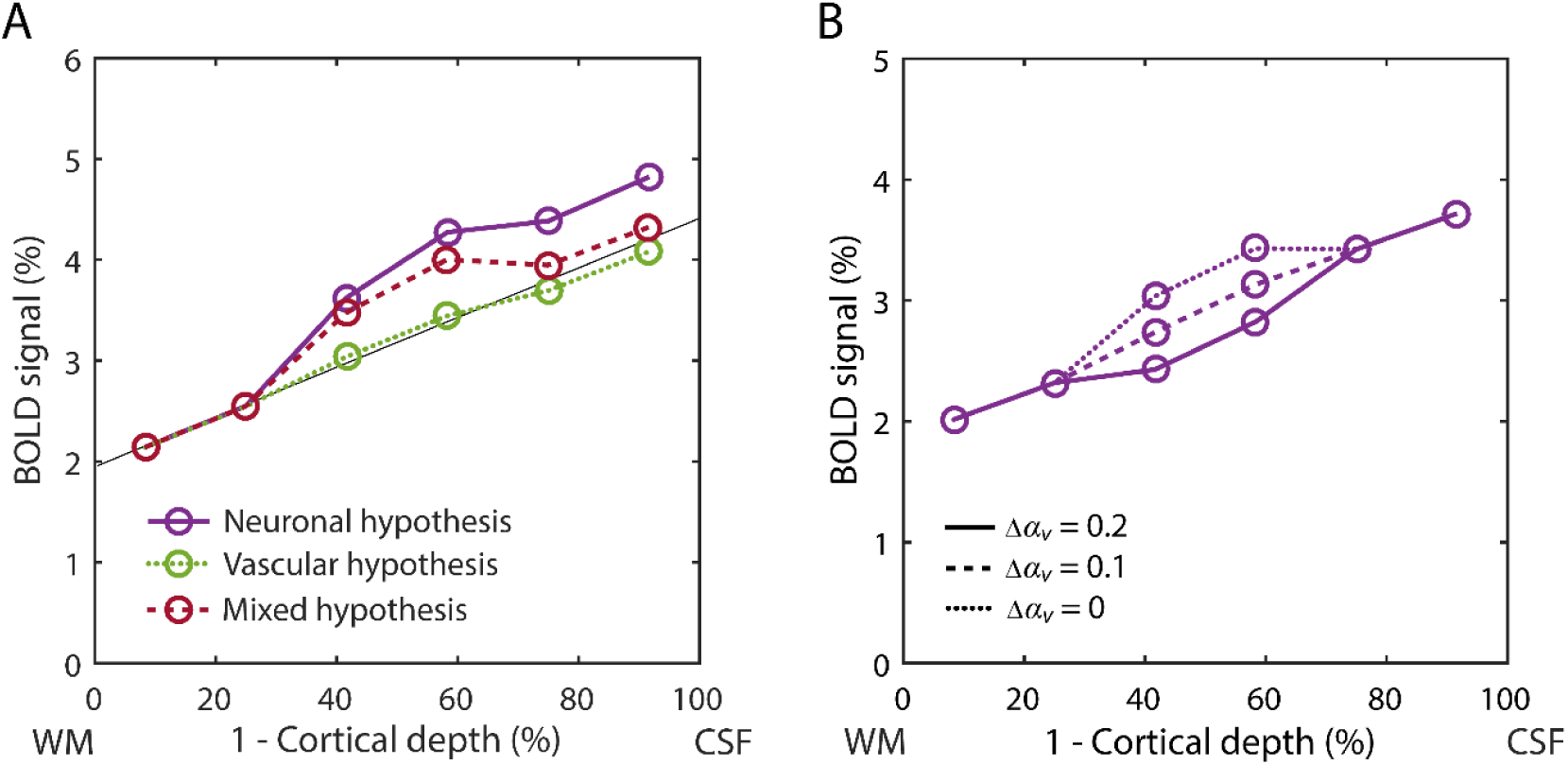
(A) Result of three simulated physiological hypotheses underlying bump in the LBR. Neuronal and mixed hypotheses show clear bump in the profile, while the bump generated by the vascular hypothesis is negligible. (B) Predicted LBRs given by different levels of laminar variation in CBF-CBV coupling. In the case of strongest variation (i.e. Δ*α*_*v*_ = 0.2), the LBR shows relative decline of signal amplitude in the middle depths, even though neuronal activity has a local maximum in the middle depths.

#### Laminar variation in CBF-CBV relationship

In Figure 5B, we show the effect of depth-specific CBF-CBV relationship on the LBR. Despite larger increase in CBF in the middle depths (by ∼10% compared to the surrounding depths), a larger CBF-CBV coupling in the middle depths (Δ*α*_*v*_ = 0.2 with respect to the surrounding depths) results in a significant decrease of the LBR amplitude in the middle depths (solid line). This is in agreement with some animal data reported by (Jin and Kim, 2008; Zhao et al., 2007a). For a smaller discrepancy in laminar CBV change (Δ*α*_*v*_ = 0.1), the effect of local maxima of CBF and *α*_*v*_ across depths almost cancels out (dashed line). For *α*_*v*_ constant across depths (Δ*α*_*v*_ = 0), we observe a considerable positive bump in the middle depths (dotted line). This corresponds to the neuronal hypothesis described above, but here simulated with a significantly smaller increase of CBF in the middle depths (i.e. 10% vs 40% considered earlier). Note that very similar dependency of LBR can be obtained if we consider variable *α*_*d*_ along the AV (data not shown).

### Results for simulation of response transients

#### Coupled vs uncoupled dynamic relationships

Figure 6 shows LBR and its underlying hemodynamic variables for dynamically *coupled* (first row) and *uncoupled* (second row) relationships between CBF & CMRO2 and CBF & CBV responses within the individual compartments of the MV and AV. In the coupled scenario, both CMRO2 and CBV response are temporally closely aligned with the CBF response in the MV and there is always a small delay between the responses in the MV and AV due to the transit time of blood. On top of this, laminar CBV and dHb responses in the AV exhibit delays, which increase towards the upper depths (see Figure 6B and C). As we will see below, the main mechanism for these intra-cortical delays is the CBV_0_ increase towards the surface in the AV. In comparison to the CBV responses, the laminar dHb responses show stronger cortical depth-dependence, with longer delays towards the surface. Additionally, laminar dHb responses contain a brief increase at the beginning of the stimulation, with its size and delay increasing towards the surface (see Figure 6C). The overall larger decrease in dHb responses in the AV is due to smaller CBV change in the MV (*α*_*v*_ = 0.35, *α*_*d*_ = 0.2). The LBRs are directly weighted by the CBV_0_. Therefore, they do show the typical increase of response peak amplitudes towards the surface (see Figure 6D). The TTP difference between lower and upper depths is ∼0.4 s. The LBR also shows an initial dip, which is related to the brief increase in dHb response, described above. This means that, in the coupled scenario, the cortical depth-dependence of the initial dip and delays in TTPs are purely the result of vascular properties (i.e. the result of CBV_0_ spatial increase towards the surface in the AV and the differences in vascular hemodynamics of the MV and AV). Finally, for the coupled scenario, the LBR does not exhibit any PSU.

**Figure 6.**
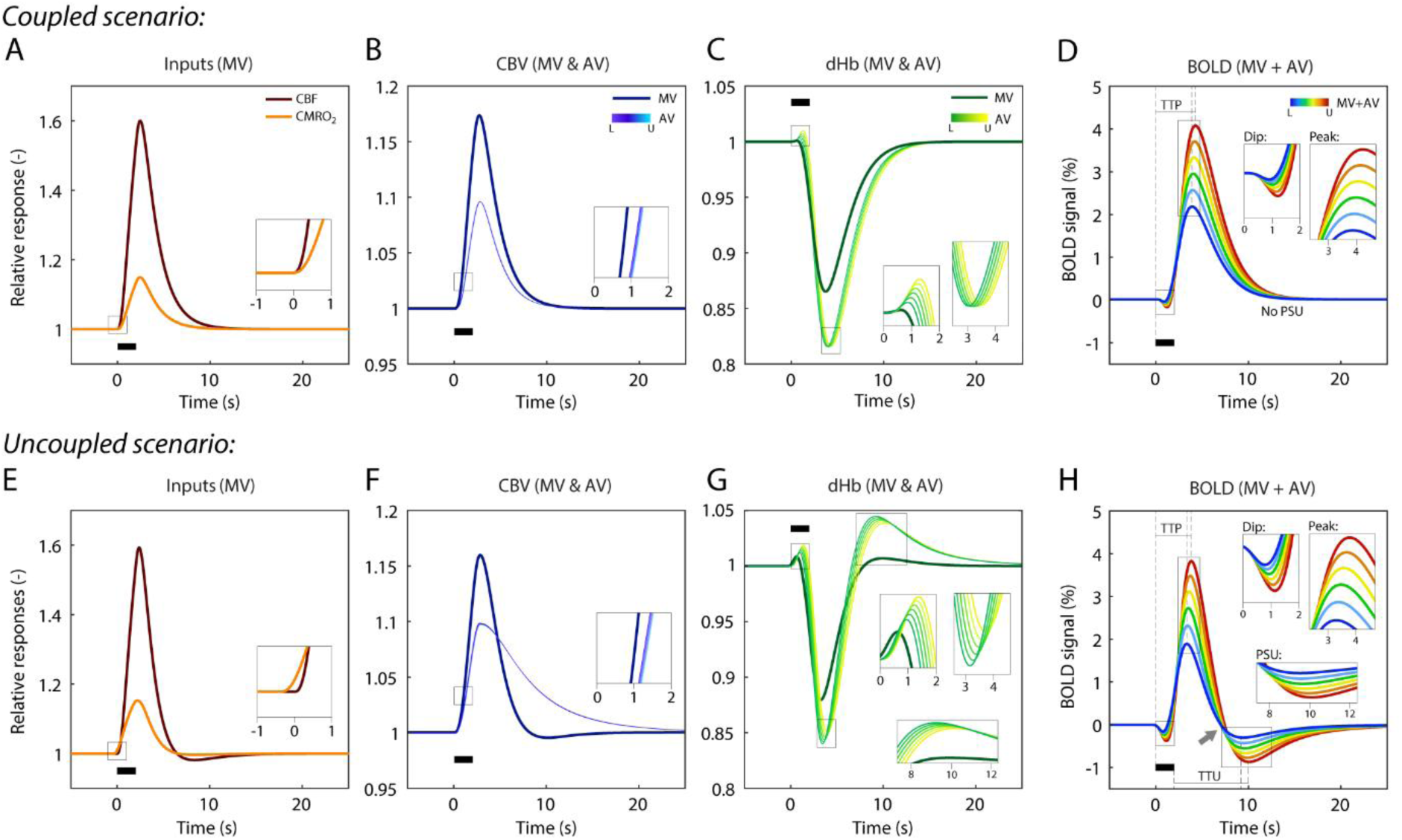
Illustration of laminar hemodynamic responses underlying LBRs during coupled (first row) and uncoupled (second row) scenarios to 2 s stimulus. Each scenario shows simulated laminar response of (from left to right): (A, E) CBF and CMRO2 in MV (i.e. model inputs); (B, F) CBV in MV and AV; (C, G) dHb in MV and AV; and (D, H) resulting LBRs as a sum of MV and AV signal components. In both scenarios, while the laminar dHb responses in MV are constant across cortical depths, laminar dHb responses in AV exhibit depth-dependence. The laminar responses between lower (L) and upper (U) depths (for AV and for LBR) are depicted using the color scale. Differences between coupled and uncoupled scenarios are emphasized with zoom-in windows; e.g. early increase of CMRO2 before CBF and post-stimulus overshoot in laminar dHb during uncoupled scenario or differences in depth-dependence of the initial dip, response peak and PSU of LBR. Note that the coupled scenario does not result in PSU of LBR (D). Measurements of TTP and TTU on LBRs are illustrated in (D and H).

Next, we simulated an uncoupled scenario with an early increase of CMRO2 and slow recovery of CBV in the AV, which induce rich dynamic features in the LBR time courses (see Figure 6F-H). In contrast to the coupled scenario, the initial dip is here predominantly due to early metabolic response. Further, the LBR includes PSU exhibiting cortical depth-dependence with TTU difference between lower and upper depths around 0.6 s. Both scenarios illustrate that the BOLD responses at different depths can strongly vary in their amplitudes and shape even if the underlying physiological parameter changes (i.e. changes in CBF, CMRO2 and CBV) are not cortical depth-specific. For quantitative evaluation of the LBR transients and how they depend on the underlying baseline and activity-induced physiological parameters, please see below and Supplementary Material 2.

#### Main factors affecting LBR transients

Spatiotemporal-dependence of LBR transients, such as early-overshoot and PSU, was further evaluated using 20 s stimulus duration. For coupled CBF-CBV changes (i.e. neuronal origin of BOLD response transients) assuming a homogenous distribution of CBV_0_ in both MV and AV results in a constant TTP and TTU^7^ across all cortical depths (see Figure 7A and B; dotted lines). Similarly, the amplitude of PSU does not exhibit any cortical depth-dependence (see Figure 7C; dotted line). Note that in the case of inhomogeneous changes in CBF or CMRO2 across depths, we observe depth-dependence of PSU amplitude (and TTP & TTU) non-locally due to signal leakage to upper depths in addition to possible local contributions (data not shown). Next, a CBV_0_ increase in AV induces a linear increase of TTP and TTU towards the surface, differing by ∼0.5 s and ∼1 s between lower and upper depths, respectively (depicted with solid lines). One can notice that while this temporal difference between lower and upper depths is comparable with TTP for short stimulus, TTU increases with longer stimulus durations. It also induces an increase of PSU amplitude towards the surface, which is relatively larger compared to the positive LBR. On the other hand, for CBV_0_ increase only in the MV (depicted with dashed lines), TTP and TTU start with much longer delays in lower depths and then strongly decrease towards the surface (i.e. resulting in about −2 s difference for TTP and −3.5 s TTU between lower and upper depths). Note that the range of TTP and TTU between lower and upper depths can also be effectively modulated by changing the CBV_0_ ratio between MV and AV. That is, larger CBV_0_ of AV results in longer inter-laminar delays of LBR transients (see Figure S2.3A in Supplementary Material 2). Additionally, changing the transit time through MV has a simple scaling effect on laminar delays and transient amplitudes (see Figure S2.3B in Supplementary Material 2).

**Figure 7.**
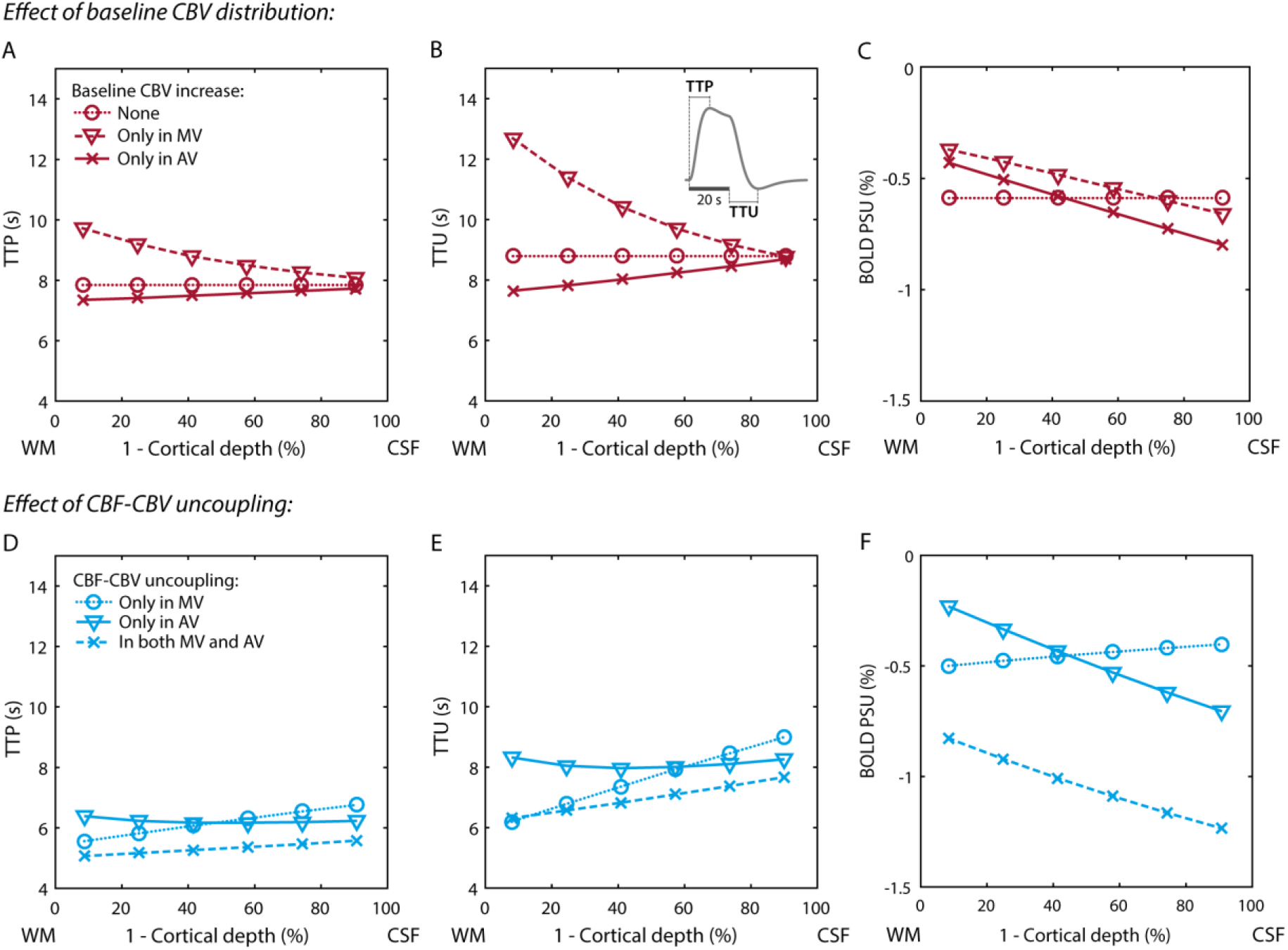
Simulation results demonstrating the effect of CBV_0_ (first row) and CBF-CBV uncoupling (second row) on cortical depth-dependence of temporal BOLD response to 20 s stimulus. Response features such as TTP, TTU and amplitude of PSU, are displayed from left to right. (A-B) CBV_0_ increase in AV (solid line) is necessary for TTP and TTU to increase towards the surface. The measurement of TTP and TTU with respect to stimulus onset and offset, respectively, is illustrated in (B). (C) CBV_0_ increase towards the surface in MV and/or AV (depicted with dashed and solid lines, respectively) is necessary for BOLD response PSU to exhibit cortical depth-dependence (i.e. PSU increasing towards the surface). (D-E) CBV-CBF uncoupling in MV or in both MV and AV results in TTP and TTU increasing towards the surface (dotted and dashed lines, respectively). CBV-CBF uncoupling only in AV (solid line) results in approximately constant TTP and TTU. (F) While CBF-CBV uncoupling only in MV leads to decrease of PSU towards the surface (dotted line), the opposite effect is achieved if the uncoupling is in the AV or in both MV and AV (solid and dashed lines, respectively).

In the case of dynamically uncoupled CBF-CBV changes, cortical depth-specific CBV_0_ (as shown above) or changes in physiological parameters are necessary for any depth dependence of LBR transients. Then, besides a significant shortening of TTP and TTU compared to the coupled relationship (by ∼2 s), CBF-CBV uncoupling can further modulate this dependence (here we consider CBV_0_ increase in the AV): If the CBF-CBV uncoupling appears only in the MV, both TTP and TTU increase towards the surface but the amplitude of PSU slightly decreases (see Figure 7D-F; dotted lines). If the CBF-CBV uncoupling appears only in the AV, both TTP and TTU remain approximately constant across depths (see Figure 7D and E; solid lines). That is, examining timing differences as a function of cortical depth can yield additional information about the location of the balloon effect (e.g. in the MV or AV). The PSU of LBR exhibits typical increase towards the surface (see Figure 7F), with the relative slope being about twice steeper compared to the positive LBR. For CBF-CBV uncoupling in both MV and AV, TTP and TTU increase towards the surface but the range between lower and upper depths (∼0.5 s and ∼1.5 s, respectively) is smaller than if the uncoupling is only in the MV (∼1.2 s and ∼2.5 s, respectively). The PSU amplitude in lower depths is larger, but the relative increase in PSU amplitude towards the surface is about twice smaller compared to the case with CBF-CBV uncoupling only in the AV (see Figure 7D-F; dashed lines).

Further, while these simulations were performed by considering six cortical depths, resulting depth dependences of BOLD response amplitudes, TTP and TTU are representative also for a different number of depths. Specifically, the depth dependence of these parameters would follow the same curve, only sampled at higher spatial resolution. It should be also clear that by increasing the number of depths the transit time through individual compartments of AV will automatically decrease. For illustration, see Figure S2.2 in Supplementary Material 2 and also Table S2 in Supplementary Material 1 indicating, which baseline physiological parameters are number of depths (in)dependent.

### Results for simulations illustrating laminar fMRI data by Huber et al. (2017)

Figure 8A and B show assumed and predicted laminar CBF and BOLD responses, respectively. Here, *x*-axis refers to time, *y*-axis to the cortical depth, and the color encodes response amplitude in percent signal change. While during the stimulation (indicated with white dashed lines), there is an increase in laminar CBF response exhibiting two peaks in lower and upper depths (indicated with green markers), the predicted positive LBR is strongly weighted towards the upper depths, with the amplitude ranging between ∼1.5% and ∼8.7%, which is in excellent agreement with experimental observations by Huber et al. (2017). Average laminar CBF and LBR over 10-30 s time-window are shown in Figure 8C. With the number of simulated cortical depths being *K* = 21, one can clearly notice that the LBR (blue line) lacks the activation-related spatial specificity simulated in the laminar CBF response (red line). That is, in the LBR, the two activation related peaks are spatially shifted towards the surface (highlighted with dashed lines in Figure 8C and with purple markers in Figure 8B) and mostly masked by the increasing slope of the LBR (for comparison with real data see Fig.2F in Huber et al. (2017)).

**Figure 8.**
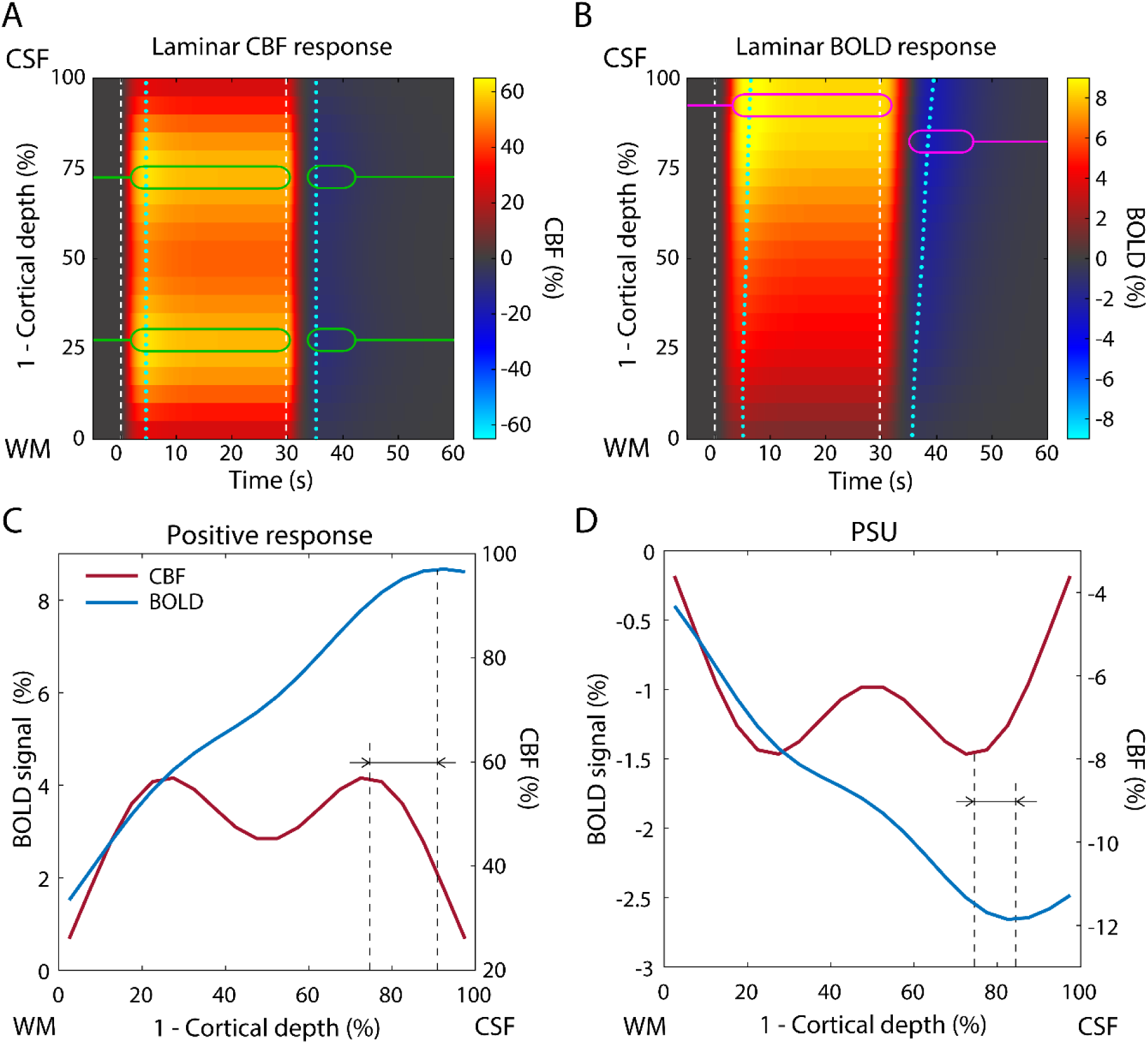
Simulation results illustrating laminar fMRI data by Huber et al. (2017) (Fig.2F). In the first row (A-B), we show activation maps of cortical depth-dependent CBF and BOLD responses to 30 s stimulus (between dashed lines). The *x*- and *y*-axes refer to time and cortical depth, respectively, with cortical surface being on the top. Green markers in the CBF map (A) highlight positions of laminar peaks in the lower and upper depths that appear during both positive response and PSU. In the BOLD map (B), shifted position of the upper peak is marked with purple markers, which differs between positive response and PSU. Position of the lower peak is not easily identifiable in the LBR due to increase of the LBR signal towards the surface. Dotted lines indicate maximal TTP and TTU across cortical depths. In the second row (C-D), we show plots of average CBF and BOLD laminar profiles during positive response (C) and PSU (D). These plots highlight differences in the spatial distribution of laminar fMRI signal across cortical depths between CBF and the BOLD signal. One can also notice difference in spatial shift of the upper peak towards the surface between the positive response and PSU (marked with dashed lines).

Next, in terms of dynamic response features, the early-overshoot and PSU in laminar CBF response are temporally aligned across depths (indicated with dotted lines in Figure 8A). On the other hand, LBR shows increasing TTP and TTU towards the surface (see Figure 8B; dotted lines). Here, because of longer stimuli (30 s) and larger CBF-CBV uncoupling (especially in the AV) compared to the simulations described above, the TTP and TTU increase more between lower and upper depths (∼1.5 s and ∼4 s, respectively). Further, Figure 8D depicts the average LBR of the PSU (averaged over 35-42 s), overlaid on the average PSU of the laminar CBF response, with two peaks (presumably due to neuronal deactivation). The BOLD response PSU does not retain laminar specificity of the CBF response but the shift of (negative) peaks is smaller compared to the positive response. This is due to lower CBF change during PSU (i.e. there is a smaller blood leakage towards the surface).

## Discussion

Here, we introduced a new dynamic laminar model of the BOLD response. This model is tailored to represent fMRI signal measured with GE sequence, which is dominated by contributions from post-capillary venous vessels (Kim and Ogawa, 2012). The total venous signal at each cortical depth is described by a local and non-local component, representing MV and AV, respectively. The model describes hemodynamics on a mesoscopic level in terms of individual cortical laminae. It is an extension of simpler macroscopic models that were developed to describe an average hemodynamic BOLD response within an ROI (Buxton et al., 1998). On the other hand, the model is a simplification of even more detailed microscopic models, such as vascular anatomical network (VAN) model (Boas et al., 2008; Gagnon et al., 2015; Markuerkiaga et al., 2016) that accounts for realistic anatomical geometry of micro- and macro-vasculature and oxygenation distribution along them. While the detailed VAN model is extremely useful for better understanding of physiological mechanisms underlying the BOLD response, it is less practical for data fitting, because it cannot be easily inverted due to very high model complexity. By saying that, simulation results obtained with the VAN model are very informative for constructing and adjusting simpler (but invertible) mesoscopic or macroscopic models.

The new laminar BOLD model is a compartmental model derived following principles of mass conservation, which allows determination of steady-state and dynamic responses. After the main physiological parameters, such as total amount CBV_0_ and CBF_0_, are chosen, the cortical depth-specific physiological parameters are unambiguously derived from these values for any number of cortical depths. Each compartment substitutes an average process within segments of the cortex; as such, it does not require the exact parameters about vessels distribution, such as their lengths and sizes, rather it is described by the average distribution of CBV_0_ and CBF_0_ divided into a specific number of cortical depths. Thus, the model retains desired level of flexibility, which ensures its generalizability across diverse anatomical and physiological conditions and will allow model inversion with a modest number of physiological assumptions in the future (see below, (Havlicek et al., 2019)). In addition, a new laminar BOLD signal equation accounting for multiple venous compartments was derived to reflect extra- and intra-vascular signal contributions and parameterized for 7 T magnetic field strength and GE-sequence. All these properties make our model also distinct from the earlier proposed (dynamic) phenomenological laminar BOLD model by Heinzle et al. (2016) that considered two-cortical depths (see below). Further, our model can be easily adapted to other field strengths and MRI sequences, such as SE, just by adjusting the BOLD signal equation (Havlicek et al., 2015; Uludağ et al., 2009). The MATLAB code provided with the publication (https://github.com/martinhavlicek/laminar_BOLD_model) allows the reader to simulate other physiological scenarios and results from high-resolution fMRI studies, not evaluated in this paper.

In principle, all physiological parameters in the model can be chosen according to available experimental data. For the forward simulations, we chose certain parameter sets, which allowed us to characterize the properties of the model or mimicked certain experimental observations in high-resolution fMRI studies (see Uludag and Blinder (2018), for recent review): (1) LBR acquired with GE sequence gradually increases from lower to upper depths (e.g. see (De Martino et al., 2013; Kashyap et al., 2017; Koopmans et al., 2010; Silva and Koretsky, 2002)); (2) Removing voxels within the cortical ribbon that contain mainly larger vessels, such as AVs, results in lowering the slope of LBR increase towards the surface and improved spatial specificity of LBR (Chen et al., 2013; Koopmans et al., 2010); (3) The depth-dependent amplitude of LBR was shown to exhibit simple amplitude scaling with stimulus intensity or contrast (Marquardt et al., 2018; van Dijk et al., 2018); (4) Studies reported relative maxima (i.e. “bumps”) in the LBR that were suggested to have either neuronal or vascular origin (Chen et al., 2013; Jin and Kim, 2008; Marquardt et al., 2018; Olman et al., 2012; Ress et al., 2007; Zhao et al., 2007a); (5) LBR transients show cortical depth-dependence as well: most commonly, the amplitudes of PSU or initial dip increase towards the surface (Kashyap et al., 2017; Tian et al., 2010; Zhao et al., 2007a); (6) LBR transients exhibit cortical depth dependent delays, which tend to increase towards the surface (Petridou and Siero, 2018; Siero et al., 2011; Tian et al., 2010). However, some studies also suggested that depth-specific variations in these delays could reflect finer mechanisms of laminar neuronal connectivity (Yu et al., 2014). Note that point (1) was earlier demonstrated using steady-state model by (Markuerkiaga et al., 2016) and points (1, 5 and 6) were also demonstrated using dynamic phenomenological model by (Heinzle et al., 2016). As another illustration and by considering model parameterization within physiological ranges, we have shown that the laminar fMRI data from Huber et al. (2017), consisting of CBV and BOLD measurements, can be remarkably well reproduced by the model – demonstrating the flexibility of the model to accommodate diverse LBR measured in animals and humans.

### Increase of LBR towards the surface

In general, the amplitudes of LBR between lower and upper depths can vary considerably as reported in several laminar studies: ranging from small (e.g. 0.5→2.5%) to large (e.g. 1.5→12%) increase towards the surface (Huber et al., 2015; Kashyap et al., 2017; Martino et al., 2013; Polimeni et al., 2010; Shen et al., 2008; Siero et al., 2011; Silva and Koretsky, 2002). We showed that under the assumption that all cortical depths are equally active (i.e. homogenous CBF activation across all depths), the typical LBR increase towards the surface can only be obtained if also the CBV_0_ spatially increases in either the MV or AV (see Figure 3A). However, a laminar increase of vascular density in the MV is not well supported by known *ex vivo* tissue observations (Weber et al., 2008), and further, it produces spatiotemporal dependence of response transients that is incompatible with most experimental data (see below for details). It is important to note that the mere draining of dHb without baseline CBV_0_ spatial increase does not result in BOLD signal increase towards the surface (see Figure 3A). This is because the transit time of dHb through AV is in this scenario much shorter in the upper depths (due to spatially increasing CBF_0_ in the AV but constant CBV_0_), which results in faster washout of dHb to the pial surface. Moreover, not considering CBV_0_ increase towards the surface together with variable CBF across depth may even result in lower amplitude of LBR in the upper compared to the lower depths (see Figure S2.3), which is inconsistent with experimental observations. Thus, in agreement with Markuerkiaga et al. (2016), our results suggest that the CBV_0_ distribution of AVs, rather than the draining of dHb alone, is the primary source of the amplitude increase of LBR towards the surface. For the same CBV_0_ distribution values as used in Markuerkiaga et al. (2016), our model closely reproduces their simulated LBRs (data not shown). Note that this result is principally different from the phenomenological laminar model by Heinzle et al. (2016). Their model does not directly distinguish between MV and AV and does not assume laminar distribution of CBV_0_. Thus, the increase of LBR amplitude towards the surface is purely the result of blood draining, parameterized with a functional coupling (i.e. parameter *λ*_*d*_ > 0) between changes in relative dHb content (and relative CBV) in lower and upper depths. On the other hand, for *λ*_*d*_ = 0 there is no blood draining and their modeled LBRs in lower and upper depths are independent. For example, a study by Chen et al. (2013) showed that removing voxels containing larger intracortical venous vessels can significantly reduce the increase of LBR amplitude towards the surface. In our model, this corresponds to reducing fraction of CBV_0_ in the AV with respect to the MV (see Figures 3B). In summary, the model is able to accommodate a broad repertoire of experimentally observed LBR that are dependent on voxel-selection and underlying vascular anatomy.

Next, we also showed that scaling the changes in relative CBF, CMRO2 or CBV results in simple (linear) scaling of the depth-dependent amplitude of LBR (see Figure 3C and Figure S2.1B and C). For example, increasing the relative CBF (or lowering relative CMRO2) scales the BOLD signal in lower and upper depths almost equally. That is, the relative slope (e.g. after normalizing to upper depth) remains the same. Simple scaling of the LBR as a function of stimulus intensity was, for example, demonstrated in studies that experimentally controlled the level of contrast or luminance of visual stimuli (Marquardt et al., 2018; van Dijk et al., 2018). On a related note, Kashyap et al. (2017) suggested that instead of using subtraction between LBR for two or more conditions, a division approach should be applied to remove the scaling factors of laminar BOLD signals. Our simulation results directly support this suggestion: For different conditions inducing various levels of neuronal activity, a deviation from linear scaling of the LBR may be a sign for a change of the spatial profile of neuronal activity. However, as we describe below, this approach does only take care of scaling factors but not of the AV draining effect.

### Laminar point-spread function: Signal leakage to upper depths

Currently, one of the most important limitations in laminar fMRI based on the GE-BOLD signal is that at one particular depth the measured response is not only influenced by signal change related to neuronal activity in that depth but also by signal from lower depths closer to WM (i.e. due to draining of dHb content by AVs towards the surface). Consequently, the observed LBRs can be remarkably inconsistent with expectations derived from electrophysiology (see also Limitations section below for a discussion of possible influence of PVs and partial volume on the laminar BOLD signal).

Therefore, we utilized the model to quantify the leakage of dHb and the resulting laminar BOLD signal PSFs during steady-state, similarly as Markuerkiaga et al. (2016), who used a different simulation approach. We demonstrated that the exact values of laminar PSFs, described by local peak and followed by tail towards the surface, are dependent on CBV_0_ distribution across depths (see Figure 4A and Figure S2.B) but they are also affected by the relative amplitude of CBF, CMRO2 or CBV changes. That is, the relative leakage to upper depths, described by the PTT ratio, considerably depends on the physiological scenario assumed (see Figure 4B). Most importantly, we showed that with increasing amplitude of relative CBF, the PTT ratio decreases (i.e. the relative leakage to upper depths increases), which means that the laminar PSFs are dependent on the level of neuronal activation. Note that in contrast to (Markuerkiaga et al., 2016), our simulation results suggest an opposite trend in PTT ratio dependence on relative CBF. At this moment, due to complex nature of the laminar hemodynamic modeling, the origin of this discrepancy is not clear. In addition, varying CBV change in lower depths (either in MV or AV) does not affect the LBR in upper depths (see Figure 5B), but the PTT is affected (due to altered BOLD signal amplitude in that specific depth). This is because the change in CBV itself does not propagate to upper depths in the AV, only blood flow and dHb *concentration* do. Note that in the phenomenological laminar model of BOLD signal by Heinzle et al. (2016), CBV change in the lower depth is designed to affect CBV change in the upper depth, which is not consistent with principles of mass conservation.

Moreover, it was suggested earlier (Markuerkiaga et al., 2016) that model-based laminar PSFs can be employed to perform deconvolution of the spatial bias in LBR. For example, Marquardt et al. (2018), relying on their steady-state simulation results, subtracted the relative influence of the lower depth from the BOLD signal of a specific depth, in order to account for the signal leakage due to AVs. Similarly, deconvolution approach based on linear regression, employing laminar PSFs as kernel functions, was also proposed by (Markuerkiaga and Norris, 2016). While our results point to possibly larger variability in derived laminar PSFs given specific physiological assumptions, deconvolution approaches mentioned-above can still be utilized but only in combination with sensitivity analysis, mapping the effect of varying model parameters within physiologically plausible ranges on estimated laminar profiles. Note that using the current model can go beyond existing approaches by directly applying nonlinear-model inversion to LBR, including both steady-state and dynamic BOLD response features.

### What is the physiological origin of the bump in the LBR?

Occasionally, a distinguishable local fMRI signal maximum (sometimes referred to as ‘bump’) is observed in the middle depths (Chen et al., 2013; Goense and Logothetis, 2006; Harel et al., 2006; Koopmans et al., 2010), deviating by ∼0.2-2% with respect to surrounding depths. Location of this bump may correlate with a higher density of vasculature around middle cortical depths, for example in the primary visual cortex (Lauwers et al., 2008; Schmid et al., 2017a; Weber et al., 2008). Therefore, this observation has led to the suggestion that the bump in the LBR may purely be a result of CBV_0_ variation across cortical depths (i.e. the vascular hypothesis). Alternatively, it is also known that the middle granular layers receive neuronal input from earlier areas in the visual system hierarchy, which can result in a larger increase of neuronal activity compared to other layers upon visual stimulation (Self et al., 2018) (i.e. neuronal hypothesis). This was directly recorded using laminar electrodes, for example in the monkey V1 region, for strong feed-forward stimuli. Please note that both scenarios are non-exclusive and may be present at the same time (i.e. mixed hypothesis).

By carefully setting up all three scenarios, we were able to show that for typical amplitudes of the physiological variables, the vascular hypothesis is unlikely, because it does only create a very small bump in the LBR. On the other hand, the neuronal hypothesis resulted in a considerable bump in the LBR that is comparable in size to experimental observations (e.g. see (Chen et al., 2013)). Further, the mixed hypothesis, which in fact may be the best representation of experimental conditions, suggested also a bump in the LBR. This scenario is comparable with simulations in Markuerkiaga et al. (2016), where a variable CBV_0_ in the MV (inspired by vascular anatomy of monkey V1 region) together with a homogenous increase in relative CBF was assumed. They could then observe that the vascular variation is reflected also in the LBR. In brief, our results suggest that the (elusive) bump is most likely due to higher neuronal activity and not a direct consequence of higher CBV_0_.

### Variable CBF-CBV coupling across depths

We also examined the effect of variable CBF-CBV coupling (i.e. variable *α*) across cortical depths, which is motivated by experimental observations in the cat visual cortex (Jin and Kim, 2008; Zhao et al., 2007a). In particular, we showed that despite higher CBF change in the middle depths, if there is proportionally even stronger CBV response in the middle depths (defined by larger *α* in the middle depths), then this can result in a relative decrease of the LBR (see Figure 5B). The spatial mismatch of local maxima in the BOLD signal and CBF or CBV corresponds very well with some experimental observations (see Fig.5A in (Jin and Kim, 2008) or Fig.5A in (Zhao et al., 2007a), and (Uludag and Blinder, 2018) for a recent review), demonstrating that relative peaks in the LBR may also not directly be associated with peaks in neuronal activity (but originating from depth-dependent variation of CBF-CBV coupling). However, one should keep in mind that the experimental data were recorded on anesthetized cats, which may not generalize to human non-anesthetized physiology (see (Uludag and Blinder, 2018), for further discussion).

### Cortical depth-dependence of BOLD response transients

The transitions between different steady-states, such as between baseline and sustained activation and *vice versa*, may be accompanied by uncoupling between different physiological variables, which can, in turn, result in BOLD response transients, such as early-stimulus overshoot, response adaptation, PSU, or initial dip. The main physiological mechanisms underlying BOLD response transients at macroscopic level were proposed to represent: (i) close reflection of changes in excitatory-inhibitory balance (i.e. neuronal origin) (Bandettini et al., 1997; Havlicek et al., 2015; Mullinger et al., 2013; Sadaghiani et al., 2009; Shmuel et al., 2006); (ii) dynamic uncoupling between CBF and CBV (i.e. vascular origin) (Buxton et al., 1998; Chen and Pike, 2009; Havlicek et al., 2017a; Mandeville et al., 1999); or (iii) dynamic uncoupling between CBF and CMRO2 (i.e. metabolic origin) (Donahue et al., 2009; Frahm et al., 2008; Lu et al., 2004; van Zijl et al., 2012). These mechanisms and, consequentially, BOLD response transients play an important role also at the mesoscopic level as their amplitudes and delays exhibit cortical depth-dependence (Petridou and Siero, 2018; Uludag and Blinder, 2018). Since the laminar BOLD model can accommodate any of the physiological mechanisms underlying the BOLD response transients (similarly as earlier introduced model by Havlicek et al. (2015)), it can provide insights into their cortical depth-dependence.

We showed (see Figure 7A and B) that it is essential to consider increasing CBV_0_ towards the surface in the AV in order to create the typically observed increase of the magnitude of the LBR transients and their delays (TTP and TTU) towards the surface (Siero et al., 2011; Tian et al., 2010). This is expected as changing CBV_0_ in the AV automatically changes the transit times through AV compartments. On the other hand, significant increase of CBV_0_ towards the surface only in the MV (as argued above to be anatomically less plausible (Weber et al., 2008)) results in decrease of TTP and TTU towards the surface (about −2 s difference for TTP and −3.5 s TTU between lower and upper depths). This is because we considered a constant transit time across depths in the MV – then assuming an increase of CBV_0_ towards the surface in the MV translates to a spatial increase in the CBF_0_ in the MV as well. Although negative trends in laminar TTP or TTU were also reported (Hirano et al., 2011; Yen et al., 2018), they appear within much smaller ranges (about −0.2 s). Thus, they are most likely due to combination of different physiological mechanisms rather than CBV_0_ spatially increasing towards the surface in the MV (e.g. variable transit time through MV across depths (Schmid et al., 2017b), see below).

Additionally, we explored the cortical depth-dependence of hemodynamic responses to short stimuli by emphasizing the difference between dynamically coupled and uncoupled relationships among different physiological variables underlying LBRs. Interestingly, for the coupled scenario we found that, even if all physiological parameters are constant across depths (such as model inputs, *α* and *n*) except CBV_0_ and CBF_0_ increasing towards the surface in the AV, the BOLD signal transients can differ in amplitude and delay up to ∼0.4 s as a function of cortical depth (see Figure 6D). Further, while the coupled scenario did not produce any BOLD response PSU, we observed a small initial dip with increasing amplitude towards the surface. This initial dip disappears for uncoupling between CBF and venous CBV or if the CBV change in the AV is negligible (see below). That is, in high-resolution fMRI, transient features can also arise even in the case of coupled physiological parameters, due to the transport of dHb through the tissue to the surface by the AVs, with the necessary condition of increasing CBV_0_ towards the surface in AVs.

Next, for uncoupling between CBF and CMRO2 during the early stimulation phase (homogenously distributed across cortical depths), the LBR showed a considerable initial dip with increasing amplitude and small delay towards the surface (see Figure 6H). This spatiotemporal dependence of the initial dip, again originating from the AV, is in excellent agreement with several studies that reported a small or negligible initial dip in lower depths but large initial dip in upper depths close to the pial surface (Siero et al., 2015; Tian et al., 2010). Note that while the same cortical dependence of the initial dip can also be created by early decrease in CBF (related to decrease in neuronal activity), this is not supported physiologically. Importantly, in either case, we observed that the amplitude increase of the initial dip towards the surface is very small if no CBV change is present in the AV. Therefore, these simulation results suggest that also some CBV change in the AV is required to create cortical depth-dependence of the initial dip with amplitudes comparable to experimental observations. That is, only an early increase in CMRO2 is not sufficient to create a significant depth-dependence of the initial dip if not accompanied by increasing CBV_0_ towards the surface and some increase of relative CBV within AVs.

The cortical depth-dependence of the BOLD response TTP was similar to the coupled scenario (i.e. increasing towards the surface) with time difference between lower and upper depths being ∼0.5 s (see Figure 6H) and remained about the same also for longer stimulus duration (see Figure 7A, solid line), which compares well with experimental studies (Hirano et al., 2011; Jin and Kim, 2008; Siero et al., 2011). During the post-stimulus period, using small PSU in the laminar CBF response and CBF-CBV uncoupling in the AV, the amplitude of LBR-PSU and TTU increased towards the cortical surface. The relative increase of PSU amplitude was larger compared to the LBR peak (or steady-state in general), which we found consistently also in simulation scenarios that assumed longer (20 s) stimulus duration (see Figure 7C and F). That is, the ratio of the amplitudes of BOLD signal during stimulation and post-stimulation cannot be directly taken to reflect relative magnitudes of neuronal activation and post-stimulus deactivation at these depths (contrary to suggestions by (Siero et al., 2015) or (Kashyap et al., 2017)). Note that significant CBF-CBV uncoupling only in MV, e.g. assuming that there is no CBV change or CBF-CBV uncoupling in the AV, results in decreasing amplitude of LBR-PSU towards the surface (see Figure 7F; dotted line), which is not commonly observed. Thus, this suggests that either, there is always some neuronal/metabolic contribution to PSU, or the CBF-CBV uncoupling takes place in both MV and AVs or mostly in AVs.

The time difference between lower and upper depths in LBR-PSU, as indicated by TTU (>1 s), was larger compared to the TTP (see Figure 6H), which is again in good agreement with experimental data (e.g. see plots with LBRs in (Siero et al., 2011; Tian et al., 2010) or (Jin and Kim, 2008)). In general, our results suggest that the inter-laminar delays in TTU increase with stimulus duration (see Figure 7B, solid line, and Figure 6H) and can be further prolonged if significant CBF-CBV uncoupling (i.e. time constant *τ* together with larger *α*) is present (see Figure 7E and Figure 8B). Note that it is commonly observed that full recovery of the BOLD response PSU to baseline can take significant amount of time (10-30 s) even for very short (e.g. 2 s) stimulus durations (Hirano et al., 2011; Hu et al., 1997). One of the most likely mechanisms behind this long recovery is the CBF-CBV uncoupling (Havlicek et al., 2017a), here parameterized with proportionally long *τ* in the MV or AV. Additionally, larger inter-laminar delays in TTP and TTU can be also obtained by increasing the fraction of CBV_0_ of the AV with respect to the MV (see Figure S2.3A). This could be related to experimental observations showing that with increasing diameter of venous vessels, hemodynamic responses exhibit longer delays (Hutchinson et al., 2006).

Thus, one should realize that underlying physiological mechanisms of hemodynamic delays in LBRs can be quite complex, resulting from interplay of possibly several physiological parameters. One should then appreciate that since our model is defined based on mass conservation law, it automatically scales for different number of depths, thus providing consistent depth-dependence of LBR amplitudes, TTP and TTU (see Figure S2.4 in Supplementary material 2). Note that also the phenomenological laminar model by (Heinzle et al., 2016) includes mechanisms to account for inter-cortical delays of LBR transients between lower and upper depths, which are in the range of typical experimental observations. In their model, these delays are controlled by draining time constant *τ*_*d*_ (which should not be mistaken for *τ*_*d*,*k*_, representing time constant of the CBF-CBV uncoupling in the AV in our model) and coupling parameter *λ*_*d*_. However, it is not very clear how these parameters generalize for a higher number of cortical depths and if they could be combined with the physiological mechanisms underlying BOLD response transients, i.e. having neuronal, metabolic or vascular origins. The standard DCM (Friston et al., 2003), which the phenomenological laminar model is based on, does not account for these important mechanisms (Havlicek et al., 2015).

In summary, as in the case of LBR amplitudes, the exact timing differences between depths of BOLD signal transients cannot be directly interpreted as timing differences of neuronal activity. Nevertheless, LBR transients, if carefully examined, offer additional means to improve the specificity of GE BOLD fMRI and could help to determine neuronal processes and neurovascular coupling relationships across cortical depths.

### Modeling BOLD response in pial veins

By extending the laminar BOLD model with a PV compartment, which collects all the drained blood from AVs, we were able to explore few basic physiological scenarios (for the blooming effect of PV, see below): first, we showed that the CBV_0_ is also the main parameter affecting the amplitude of PV response at the cortical surface (see Figure A1). For considerably larger CBV_0_ of PV than AV, the amplitude of PV signal increases following the LBR. For CBV_0_ of PV that is the same or less than CBV_0_ of AV, the amplitude of PV signal is lower than the LBR in the upper depth. Both cases were experimentally observed (Ress et al., 2007; Siero et al., 2011; Siero et al., 2013).

Following the uncoupled scenario described earlier, we also examined BOLD response transients in the PV (Figure A1). Altered dHb *concentration* in the AV propagates to the PVs and therefore evokes BOLD signal transients that are non-local in origin (see Figure A1, blue line). This means that one can, for example, observe in the large surface vessels (i.e. in voxels containing PVs) a BOLD response with PSU, even without slower return of CBV to baseline in the PV, as reported by Yacoub et al. (2006). In this case, we expect that the ratio of PSU to positive BOLD response is much smaller in the PV compared to upper depths of GM. In other words, if the observed ratio of PSU to positive BOLD response is comparable or larger than the ratio in the upper depths of GM, it is an indication that additionally, the CBF-CBV uncoupling is contributing to BOLD response PSU in the PV (see Figure A1, orange line). Further, TTP and TTU of PV response are longer compared to LBRs within GM. All these simulation results are plausible and in agreement with experimental observations (e.g. see (Petridou and Siero, 2018; Siero et al., 2013; Tian et al., 2010)).

### Limitations and future prospects

- The new model of the LBR is based on modeling hemodynamic changes in venous compartments (similarly as the original balloon model (Buxton et al., 1998)). This is because the fMRI BOLD signal acquired at 7 T with GE sequence is dominated by the fMRI signals coming from and around venous vessels. Theoretical simulations based on realistic VAN model (Boas et al., 2008; Gagnon et al., 2015) clearly showed that venous vessels contribute by ∼80%, capillaries by ∼20% and contribution from arterial vessels is close to zero. These results are well aligned with many other theoretical studies (e.g. see (Kim and Ogawa, 2012; Ogawa et al., 1993; Uludağ et al., 2009). Although it is known, especially from optical imaging studies, that there are larger CBV changes at the arterial side (in particular for short stimulus durations) (Hillman et al., 2007; Vazquez et al., 2010), the CBV_0_ of arterial vessels is less than 1/3 compared to that in the capillaries and venous vessels (Gagnon et al., 2015; Schmid et al., 2017a), which together with small amount of dHb in arterial vessels makes the arterial contribution to the fMRI BOLD signal small, and, thus, can be ignored to capture the main effects influencing the LBR. By saying that, our depth-specific model compartments of MV do not represent only venules, but rather a simplified model of both capillaries and venules, being more weighted towards venules. Note that the input to our model is CBF in arterioles, which can be directly linked to CBF measured with ASL data (Havlicek et al., 2017c). However, for small effect sizes (such as the amplitude of the initial dip (Uludağ, 2010)) or reduced oxygenation values in arteries (e.g. for hypoxia), the contribution of arteries and arterioles to the fMRI signal may be relevant. Additional vascular compartments can, however, be straightforwardly included in the laminar BOLD model albeit by introducing more parameters and increasing its complexity.
- In the current paper, we have set up the laminar BOLD model for 7 T and a GE acquisition (see Table 2), as these are still the most widely used MR field strength and contrast mechanism for laminar fMRI of the human brain. However, our model can be easily adjusted also for different magnetic field strength or other BOLD-sensitive sequences (Havlicek et al., 2015; Uludağ et al., 2009), by reparametrizing the BOLD signal equation (6). That is, the physiological part of the model is independent of the specific magnetic field strength and sequence parameters. Furthermore, the extension of the laminar BOLD model to multimodal fMRI data that measure total CBV (with VASO sequence) or CBF (with arterial spin labeling (ASL) sequence (Liu and Brown, 2007)) next to the BOLD signal is straightforward but beyond the scope of the current paper.
- The model does not account for effects related to voxel size and its orientation with respect to cortex curvature, generally referred to as partial volume effect (PVE). The PVE was considered in the steady-state model by Markuerkiaga et al. (2016) and it results in additional blurring of the laminar profile. It was suggested earlier that for a given voxel size, a convolution kernel can be derived (Koopmans et al., 2011) and then be applied to simulated laminar data. For actual fitting of model predictions to real fMRI laminar data, modeling of PVE is desirable. In that case, one could also utilize the PV compartment on the cortical surface (see Appendix); i.e. to model the PVE between PV and GM voxels. Additionally, LBR amplitude is dependent on blood vessel orientation with respect to the external magnetic field (B0). While this is not much of concern for MV, for which vessel’s orientation is mostly random, the AVs are oriented perpendicularly to the surface, which, if oriented 90° with respect to B0, can result in significant signal reduction (Fracasso et al., 2018; Gagnon et al., 2015). Accounting for this effect is more relevant for a small ROI with high variation in cortex curvature. In that case, it is possible to extend/modify the BOLD signal equation (6) to account for this effect.
- The magnetic field disturbances due to susceptibility changes in PVs can extend to remote tissue voxels (the so-called blooming effect, see e.g. (Kashyap et al., 2018)). This can result in additional fMRI signal changes in the GM voxels in close proximity to PVs (see, for example, recent reports of (Kay et al., 2019; Moerel et al., 2018)). Although we model dHb and CBV changes in extra- and intra-vascular compartments of PVs, the contribution of the blooming effect to the laminar BOLD signal is not accounted for in the proposed model. In order to do this from an MRI physics point-of-view, first, the susceptibility changes in PVs must be determined close to the ROI being investigated and, second, the field distribution (as a function of distance and orientation) and the ensuing fMRI signal must be estimated (Martindale et al., 2008; Ogawa et al., 1993).
- Many physiological parameters can in principle vary across cortical depths. Motivated by experimental observations, our simulations considered: (1) Laminar variations in CBV_0_ and CBF_0_ with corresponding variations of depth-specific transit times in the AV; and (2) Laminar variations in relative CBF changes and in CBF-CBV relationship. While in this paper, we kept other parameters constant across cortical depths, practically speaking, they all can be defined as depth-specific. For example, experimental data show that there may be depth-specific distribution of baseline 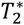 in the GM. This variation is likely due to variable iron/myelin concentration. The laminar BOLD model is derived from the baseline signal conditions. Thus, it is straightforward to apply depth-specific values of 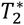, as reported e.g. by (Koopmans et al., 2011; Zhao et al., 2006). Our simulations accounting for this variation showed a negligible effect on the LBR (results not shown). Further, one can also consider depth-specific transit times through the MV (*t*_*v*0,*k*_). Although transit time through capillaries should not be too heterogenous across depths, as this results in inefficient oxygen extraction (Jespersen and Østergaard, 2012), recent experimental and theoretical findings (Gutierrez-Jimenez et al., 2016; Schmid et al., 2017b) suggest that capillary transit time can increase with cortical depth. This means that also the baseline oxygen extraction fraction *E*_0_ may be slightly different between cortical depths both in the MV and AV (Li et al., 2019). While in the current implementation one can define depth-specific changes in oxygen metabolism (directly with *m*_*k*_(*t*) or via ratio *n*_*k*_), *E*_0,*k*_ can be specified within laminar BOLD signal equations (see derivation details in Supplementary Material 1). Thus, with the MATLAB code provided with the publication, we encourage the readers to explore these and other biophysical scenarios.
- Our model is designed to be used in combination with a laminar neuronal model to estimate the spatiotemporal profile of neuronal activity using the Bayesian estimation framework (Friston et al., 2007). While in this work, we aimed for providing a general description of the model and its parameters, future work will focus on model complexity (i.e. number of free parameters) and separability of neuronal from vascular parameters in order to make the model invertible. To this end and despite above-mentioned plausibility of depth-specific variations in many physiological parameters, we foresee that it may be sufficient for most of the model parameters to be considered constant across depths. This means that the complexity of the laminar hemodynamic model does not necessarily need to increase with the increasing number of cortical depths. Even the increase of CBV_0_ towards the surface in the AV, which is one of the essential parameters to describe spatiotemporal properties of LBRs, can be controlled by a single parameter (*s*_*d*_). Further, sufficient variation in depth-dependent LBR response transients can be modeled by varying steady-state and dynamic CBF-CBV relationships (*α*_*d*_ and *τ*_*d*_) in the AV and fraction of CBV_0_ in the AV (1 − *ω*_*v*_) with respect to the MV. In other words, it is likely that for model inversion, we can retain physiological realism of our laminar hemodynamic model defined, for example, for six depths with a number of free parameters that compares to the number of free model parameters in the phenomenological laminar model by (Heinzle et al., 2016) defined for two depths. On the other hand, a higher number of free parameters can be utilized for the laminar neuronal model. Practically speaking, the number of free model parameters, which can be determined through model inversion, will depend on the quality and complexity of the experimental laminar fMRI data. Therefore, utilizing experimental manipulations that can elicit changes in LBRs together with modeling dynamic transients of the LBR will be extremely important for successful model inversion. Recently, we demonstrated with preliminary results based on simulated data (Havlicek et al., 2019) that, with the temporal constraints given by experimental manipulations (i.e. unique laminar profiles induced by different conditions) and spatiotemporal constraints given by the hemodynamic model of laminar BOLD signal, model inversion and estimation of the underlying neuronal activity is feasible.

## Summary

In summary, we propose a new dynamic generative model of the laminar BOLD response, which takes into account the effect of intra-cortical ascending veins. We illustrated the versatility of the proposed model to characterize common experimental observations in laminar fMRI: (1) We showed that the spatial increase of LBR towards the pial surface is mainly due to CBV_0_ and less due to drainage of dHb; (2) Local variability in laminar BOLD profile (i.e. bumps) is more likely due to variability in neuronal activity (i.e. ensuing CBF) rather than locally higher CBV_0_, but it can also be affected by variability in CBF-CBV (or CBF-CMRO2) coupling across depths; (3) We showed that, by assuming different physiological scenarios, the model can be used to simulate cortical depth-dependence of transient features, such as early-overshoot, post-stimulus undershoot, or initial dip. Finally, the proposed model is fully scalable to an arbitrary number of cortical depths and can be used in the future as a tool to infer spatiotemporal distributions of neuronal activity from experimentally controlled laminar fMRI data using Bayesian model inversion.

## Supporting information

Supplementary Material 1 (Model derivation)

Supplementary Material 2 (Additional simulations and results)

## Acknowledgments

We would like to thank to Drs. Sriranga Kashyap for helpful comments on an earlier version of the manuscript. This work was financially supported by a VIDI grant (#452-11-002) of the Netherlands Organization for Scientific Research (NWO) and funding from IBS (#IBS-R015-D1) to KU. MH was also partly financed by NIH grant (#RF1MH116978).

## Appendix

In the Methods section, we described the dynamic physiological states in terms of absolute variables and defined their baseline values (see also Supplementary Material 1). For simulation purposes, it is more straightforward to use normalized variables and use absolute baseline values as scaling factors. Thus, the relative variables were obtained by normalizing the absolute variables with respect to their baseline values: *f*_*i*,*k*_(*t*) = *F*_*i*,*k*_(*t*)/*F*_0*i*,*k*_, *q*_*i*,*k*_(*t*) = *Q*_*i*,*k*_(*t*)/*Q*_0*i*,*k*_, and *v*_*i*,*k*_(*t*) = *V*_*i*,*k*_(*t*)/*V*_0*i*,*k*_. After substituting the absolute variables in Equations (1-2) by their relative counterparts and further rearrangements (see Supplementary Material 1 for detail derivation), we obtain:

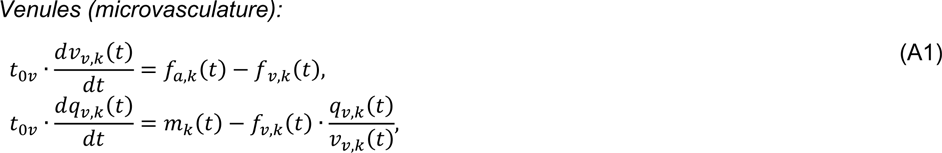

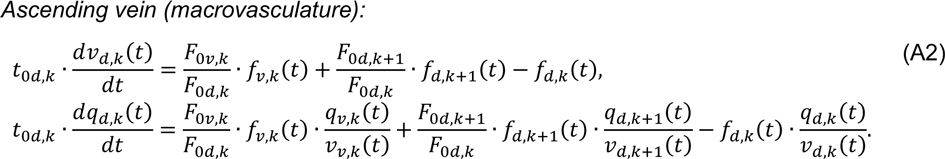

One can notice that the mass balance equations for the venules compartments have the same form as in the standard balloon model (Buxton et al., 2004; Buxton et al., 1998). Here, *t*_0*v*,*k*_ = *V*_0*v*,*k*_/*F*_0*v*,*k*_ are the mean transit times for blood to pass through the MV. Further, *m*_*k*_(*t*) represents the relative change in CMRO2, generally defined as: *m*_*k*_(*t*) = *E*_*k*_(*t*)/*E*_0_ · *f*_*a*,*k*_(*t*), where *E*_0_ is the baseline oxygen extraction fraction. Under the assumption of linear relationship between blood flow and oxygen metabolism parametrized by *n*-ratio at steady-state (Buxton et al., 2004) (considered here constant across depths), this transforms to *m*_*k*_(*t*) = (*f*_*a*,*k*_(*t*) + *n*_*k*_ − 1)/*n*_*k*_ (Buxton et al., 2004). Alternately, one can define *m*_*k*_(*t*) to be independent and uncoupled of blood flow as the second driving function of the model (Buxton et al., 2004; Obata et al., 2004).

In the AV (Equation A2), the relative changes in CBV and dHb are also scaled by the mean transit times through individual compartments, which are calculated directly from distributions of baseline CBV and CBF in the AV, *t*_0*d*,*k*_ = *V*_0*d*,*k*_/*F*_0*d*,*k*_. Additionally, the blood inflows are further scaled by ratios of baseline CBFs from different compartments. This is a result of mass conservation law due to merging of two (possibly different) baseline blood flows into a single compartment (for details see Supplementary Material 1). One can notice that the relative dHb concentrations, multiplied with corresponding relative blood flows, are passed between compartments. That is, at *k*-th depth, the AV compartment collects the dHb concentrations and flows from the MV at the same depth, *q*_*v*,*k*_(*t*)/*v*_*v*,*k*_(*t*) · *f*_*v*,*k*_(*t*), and from the lower depth of the AV, *q*_*d*,*k*+1_(*t*)/*v*_*d*,*k*+1_(*t*) · ^*f*^*d*,*k*+1(*t*).

### Pial vein (macrovasculature)

Here we also describe relative CBV and dHb changes in the PV, which similarly as the standard balloon model is represented by a single vascular compartment:

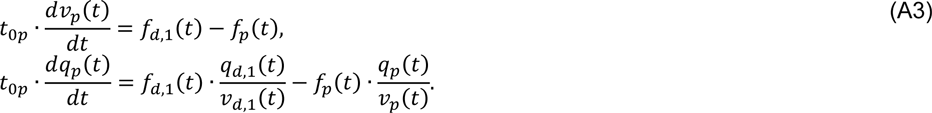

The mean transit time trough PV equals the ratio of CBV_0_ and CBF_0_, *t*_0*p*_ = *V*_0*p*_/*F*_0*p*_. The CBV_0_ in PV, *V*_0*p*_, is defined independently of the total amount of venous CBV_0_ within GM, *V*_0_, and the CBF_0_ in PV equals CBF_0_ leaving the AV, which also equals the total amount of CBF_0_ delivered by arteries, *F*_0*p*_ = *F*_0*d*,1_ = *F*_0_. The inputs to the PV compartment are flow and dHb concentration coming from the most superficial depth of AV (see also Figure 1). As for MV and AV, the CBF-CBV relationship in the PV is defined with (Equation 6). Since the PV compartment is considered as an additional zeroth-depth, it has its own BOLD signal equation consisting of extra- and intra-vascular signal contributions and related constants:

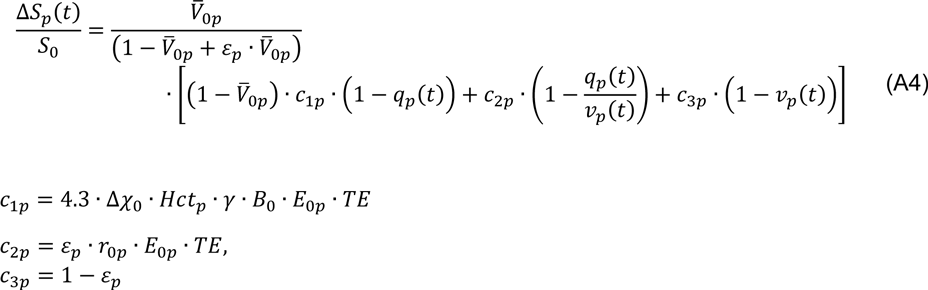

This equation has a very similar form as derived by (Stephan et al., 2007). The only differences are related to the scaling constants. In Stephan et al., the following approximations are applied during derivation: denominator 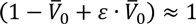 and ratio 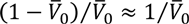. Detailed derivation is included in the Supplementary Material 1. Note that possible effects of CSF contributions are ignored as its dynamic response is expected to have a very comparable shape as the BOLD response of PV surrounded completely by GM.

In Figure A1, we illustrate two examples of the PV response time-course. These simulations of PV BOLD response followed the hemodynamic response for MV and AV designed for the uncoupled scenario, as described in Figure 6E-H. The first example shows the case when CBV_0_ of PV is half of the total amount of CBV_0_ within GM (i.e. MV+AV) and with no CBV change in the PV. This results in the BOLD response peak amplitude of PV to be smaller compared to the upper depth of GM. The TTP of the PV BOLD response further increased compared to the upper depth of GM (by ∼0.25 s). In terms of response transients, the initial dip is negligible and the PSU of BOLD response is small. Note that the effect of CBF-CBV uncoupling does not propagate between compartments (i.e. viscoelasticity is a local property of the vascular compartment). Thus, even though there was a significant CBF-CBV uncoupling in the AV, it has no effect on the PSU of BOLD response in the PV. Here it is caused by the post-stimulus CBF decrease that originated in the MV. The ratio between PSU and positive BOLD response in PV is ∼0.05, which is ∼4.2 times lower than the BOLD response in the superficial depth (∼0.21). The TTU in the PV is significantly longer compared to the upper depth of GM (by ∼1.5 s).

**Figure A1.**
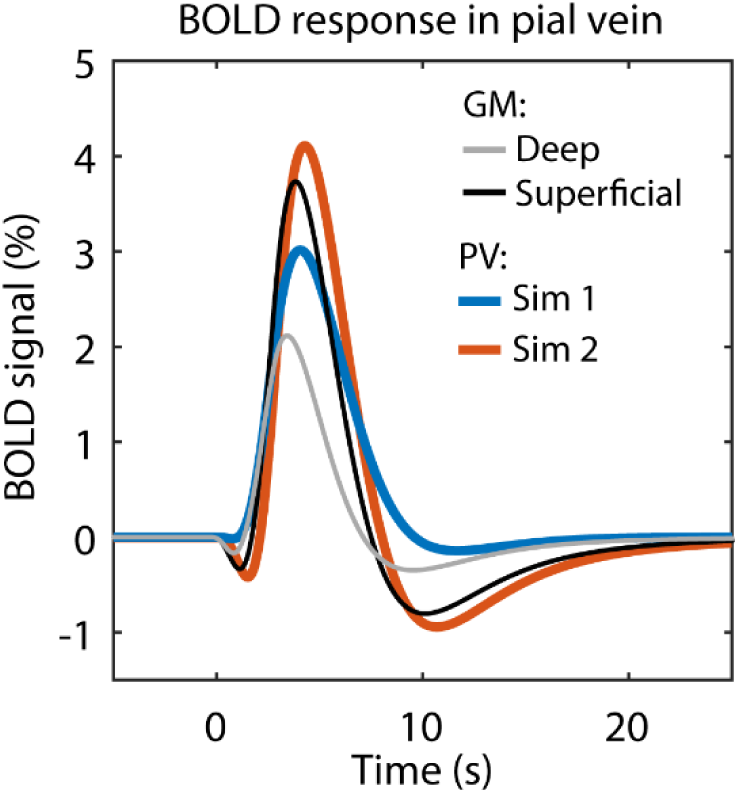
Simulated BOLD response in the PV. This plot depicts two simulation scenarios Sim 1 and Sim 2, both following simulation results of the uncoupled scenario in Figure 6H for GM (here only deep and superficial depths are shown): Sim 1 (blue line) assumes CBV_0_ of PV to be half of the AV and no CBV change; Sim 2 (orange line) assumes the same CBV_0_ as in the AV and also some CBV change and CBF-CBV uncoupling (*α*_*p*_ = 0.2, *τ*_*p*+_ = 2 s and *τ*_*p*−_ = 20 s).

The second example considers CBV_0_ in the PV to be equal to the total CBV_0_ within GM, and the same CBV change and CBF-CBV uncoupling as in the AV are assumed. Then, the PV BOLD response peak is larger compared to the upper depth of GM (still reduced due to CBV change). Larger CBV_0_ in the PV increased the TTP (by additional ∼0.2 s). Both response transients are more pronounced: The initial dip increased due to relatively tight CBV-CBV coupling in the PV during inflation phase. The size of BOLD response PSU significantly increased due to CBV-CBV coupling during deflation phase. The ratio between PSU and positive BOLD response in PV is ∼0.23, which is only slightly more compared to the BOLD response in the superficial depth. Finally, the TTU is longer than in the upper depth of GM (by ∼0.6 s) but shorter compared to the first example.

---

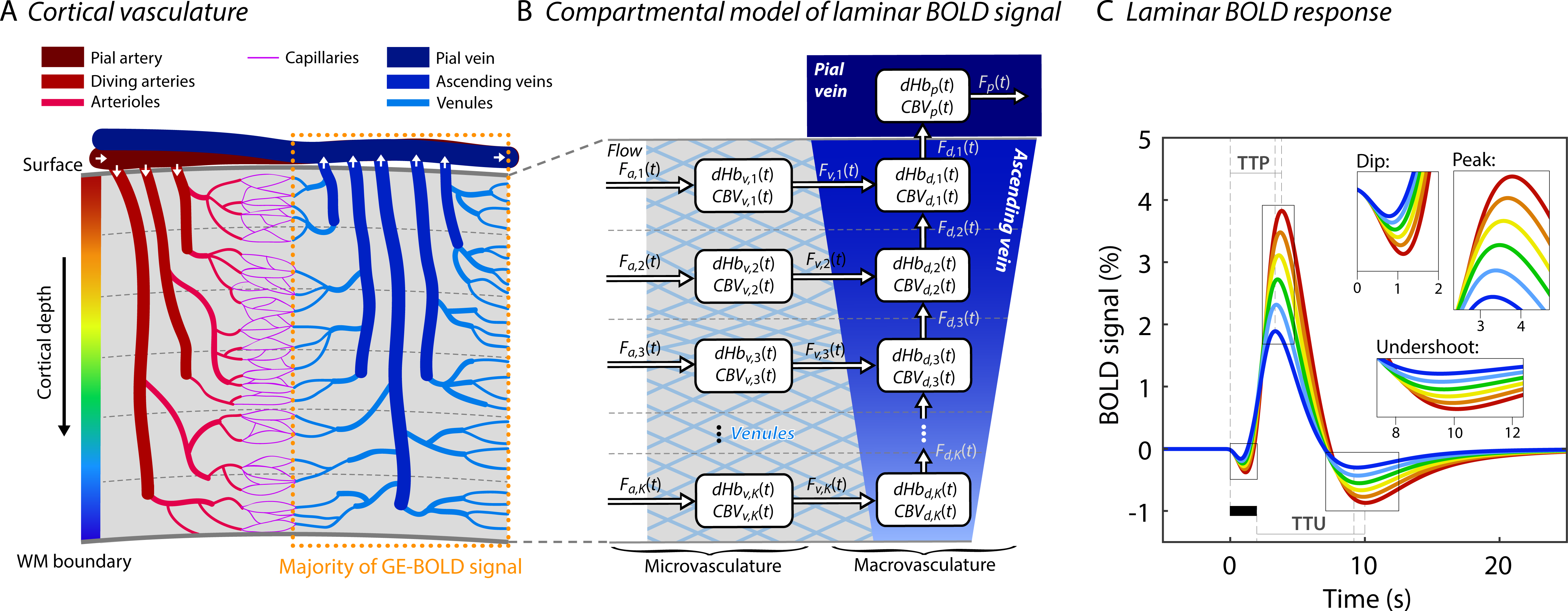

## Footnotes

1 See Table 1 for list of used abbreviations.

2 In order to avoid confusion with histological layers, we use the term cortical depth for the vertical division (i.e. pial to white matter) of the cortical tissue. The number of cortical depths can be arbitrarily chosen, whereas the number of histological layers is usually cytoarchitectonically defined – typically six layers in the human neocortex.

3 In order to distinguish from the temporal properties of the BOLD signal and from the BOLD response at a single depth, in the following we use LBR for the entire spatial profile of the BOLD signal as a function of cortical depth.

4 Note that the taxonomy of cortical blood vessels varies in the literature. In this paper, we adhere to the most commonly used terms within the fMRI literature (Uludag and Blinder (2018); Petridou and Siero (2018)). For example, some studies consider DAs and AVs as arterioles and venules, respectively. Here we reserve the terms arterioles and venules only for microvasculature and prefer to use the term vein for macrovasculature, i.e. for ascending and pial veins.

5 Note that in MRI and PET literature, CBF reflects blood flow in arterioles and capillaries, while here for simplicity we use the term also to describe blood flow in venous compartments. That is, we distinguish between CBF in the arterioles, venules, AV and PV. Further, we assumed that the CBF_0_ in arterioles equals the CBF_0_ in venules and the average CBF_0_ in capillaries (see Figure 1D).

6 Note that this is a reasonable assumption, since we expect that the laminar profile of the relative CBV and CBV in absolute units are very similar due to relatively homogenous distribution of baseline CBV in the MV. Further, for this type of experimental stimulation, it is common to observe a relative change in total CBV between 10 to 30% (Chen and Pike, 2009), which by assuming *α* between 0.4 and 0.5 (larger CBV changes are expected on arterial side) can be converted to relative CBF change between 30 to 60%.

7 Note that, we focus our description more on the time differences in TTP and TTU between upper and lower depths, and less on the actual TTP or TTU in the BOLD response because they are strongly dependent on the assumed TTP and TTU in the CBF response and transit time through the MV.

