## Supplementary Material 1 (Model derivation) for "A dynamical model of the laminar BOLD response"

### 1. Derivation of laminar hemodynamic model

Below we provide a step-by-step derivation of the laminar hemodynamic model. We start from a more general equation of mass balance:

$$\frac{d(C(t)\cdot V(t))}{dt} = C_{in}(t)\cdot F_{in}(t) - C(t)\cdot F_{out}(t) + \frac{d\left(C_{prod}(t)\cdot V(t)\right)}{dt} \tag{S1.1}$$

This equation says that a mass accumulation rate equals the mass flux into the compartment minus the mass flux out of the compartment, plus the rate of mass production. Here, the mass accumulation rate is given by a product of concentration $C(t)$ and volume $V(t)$. The mass flux is described by a product of concentration and volumetric flow rate $F(t)$[1]. Further, since we assume well-mixed compartment, the concentration leaving the compartment is equal to the concertation inside the compartment $C(t)$. The mass production rate in Equation S1.1 can be neglected in our case because we are interested in modeling dHb changes in venous vessels where no further oxygen extraction is expected. However, we do expect that blood vessels can change their volume with time. Thus, since dHb concentration equals $C(t) = Q(t)/V(t)$, we can express the mass balance equation in terms of absolute changes in dHb content, $Q(t)$, and volume, $V(t)$, as:

$$\begin{aligned} \frac{dQ(t)}{dt} &= C_{in}(t)\cdot F_{in}(t) - C(t)\cdot F_{out}(t) \\ \frac{dV(t)}{dt} &= F_{in}(t) - F_{out}(t) \end{aligned} \tag{S1.2}$$

In the following, we formulate these mass balance equations as depth-specific for different vascular compartments including venules, AV and PV. We start by defining each compartment in terms of absolute variables (as above, expressed with capital letters) and then reformulate them in terms of relative variables (using lowercase letters) with respect to baseline values (using capital letters with 0 subscript). Below, we use subscript $k$ to denote depth with respect

[1] Note that volumetric flow rate can be related to blood velocity $v$ via $F = v\cdot A$, where $A$ is the cross-sectional area, or to pressure $P$ and blood vessel resistance $R$ via $F = \Delta P/R$, where $\Delta P$ is pressure difference between two ends of vessel. The vessel resistance is further dependent on vessel diameter and length.

to the cortical surface, $a$ for arterioles, $v$ for venules, $d$ for AV (referring to draining effect), and $p$ for PV.

*Venules compartments:*

Similarly as above, dynamic changes in dHb content within venules compartment are described by dHb flux entering and leaving the venules compartments:

$$\frac{dQ_{v,k}(t)}{dt} = F_{a,k}(t) \cdot E_k(t) \cdot C_{a,k}(t) - F_{v,k}(t) \cdot C_{v,k}(t) \tag{S1.3}$$

The dHb flux entering the compartment is given by volumetric flow rate in arterioles $F_{a,k}(t)$ times oxygen extraction fraction $E_k(t)$, describing amount of oxygen extracted by capillaries, times oxygen concertation in arterioles $C_{a,k}(t)$ (for simplicity we assume fully oxygenated blood). This flux also corresponds to cerebral metabolic rate of oxygen (CMRO$_2$), $M_k(t) = F_{a,k}(t) \cdot E_k(t) \cdot C_{a,k}(t)$.

Next, normalizing Equation S1.3 with respect to baseline values yields:

$$\begin{gathered}\frac{dQ_{v,k}(t)/Q_{0v,k}}{dt} = \frac{F_{0v,k} \cdot C_{0v,k}}{Q_{0v,k}} \cdot \left[\frac{F_{a,k}(t)}{F_{0v,k}} \cdot E_k(t) \cdot \frac{C_{a,k}(t)}{C_{0v,k}} - \frac{F_{v,k}(t)}{F_{0v,k}} \cdot \frac{C_{v,k}(t)}{C_{0v,k}}\right], \\ \frac{dq_{v,k}(t)}{dt} = \frac{F_{0v,k}}{V_{0v,k}} \cdot \left[f_{a,k}(t) \cdot \frac{E_k(t)}{E_{0,k}} - f_{v,k}(t) \cdot \frac{q_{v,k}(t)}{v_{v,k}(t)}\right],\end{gathered} \tag{S1.4}$$

$C_{0v,k}$ is the baseline concentration of dHb in the venules, which is given by $C_{0v,k} = E_{0,k} \cdot C_{0a,k}$, and because we assume fully oxygenated blood in arterioles, $C_{a,k}(t)/C_{0a,k} = 1$. During baseline, the flow rates entering and leaving venules compartment are the same $F_{0v,k} = F_{0a,k}$, thus $f_{a,k}(t) = F_{a,k}(t)/F_{0v,k}$. Additionally, we make use of general relationships:

$$\frac{F_0 C_0}{Q_0} = \frac{F_0}{V_0} = \frac{1}{t_0}, \tag{S1.5}$$

$$c(t) = \frac{C(t)}{C_0} = \frac{Q(t)}{V(t)} \cdot \frac{V_0}{Q_0} = \frac{q(t)}{v(t)}. \tag{S1.6}$$

Putting everything together yields:

$$t_{0v,k} \cdot \frac{dq_{v,k}(t)}{dt} = f_{a,k}(t) \cdot \frac{E_k(t)}{E_{0,k}} - f_{v,k}(t) \cdot \frac{q_{v,k}(t)}{v_{v,k}(t)}, \tag{S1.7}$$

Since $m_k(t) = M_k(t)/M_{0,k}(t) = f_{a,k}(t) \cdot E_k(t)/E_{0,k}$, then:

$$t_{0v,k} \cdot \frac{dq_{v,k}(t)}{dt} = m_k(t) - f_{v,k}(t) \cdot \frac{q_{v,k}(t)}{v_{v,k}(t)}, \tag{S1.8}$$

For CBV changes in venules, derivation to obtain relative changes yields:

$$
\begin{aligned}
\frac{dV_{v,k}(t)}{dt} &= F_{a,k}(t) - F_{v,k}(t), \\
\frac{dV_{v,k}(t)/V_{0v,k}}{dt} &= \frac{F_{0v,k}}{V_{0v,k}} \cdot \left[\frac{F_{a,k}(t)}{F_{0v,k}} - \frac{F_{v,k}(t)}{F_{0v,k}}\right], \\
t_{0v,k} \cdot \frac{dv_{v,k}(t)}{dt} &= f_{a,k}(t) - f_{v,k}(t).
\end{aligned}
\tag{S1.9}
$$

*Ascending vein compartments:*

Within AV at specific depth $k$, dynamic changes in dHb content are described by dHb flux entering from venules at the same depth, dHb flux entering from the lower depth of AV and dHb flux leaving the AV compartment:

$$
\begin{aligned}
\frac{dQ_{d,k}(t)}{dt} &= F_{v,k}(t) \cdot C_{v,k}(t) + F_{d,k+1}(t) \cdot C_{d,k+1}(t) - F_{d,k}(t) \cdot C_{d,k}(t) \\
\frac{dQ_{d,k}(t)/Q_{0d,k}}{dt} &= \frac{C_{0d,k}}{Q_{0d,k}} \cdot \left[\frac{F_{0v,k} \cdot F_{v,k}(t)}{F_{0v,k}} \cdot \frac{C_{v,k}(t)}{C_{0d,k}}\right. \\
&\qquad \left. + \frac{F_{0d,k+1} \cdot F_{d,k+1}(t)}{F_{0d,k+1}} \cdot \frac{C_{d,k+1}(t)}{C_{0d,k}} - \frac{F_{0d,k} \cdot F_{d,k}(t)}{F_{0d,k}} \cdot \frac{C_{d,k}(t)}{C_{0d,k}}\right].
\end{aligned}
\tag{S1.10}
$$

Now, because we assume that there is no further oxygen extraction in the within venous vessels, the baseline dHb concentrations equal between all venous comportments:

$$
C_{0v,k} = C_{0d,k} = C_{0d,k+1}. \tag{S1.11}
$$

With this assumption, the relative change in dHb content within AV compartment at $k$-depth is given by:

$$
\begin{aligned}
\frac{dq_{d,k}(t)}{dt} &= \frac{C_{0d,k} \cdot F_{0d,k}}{Q_{0d,k}} \cdot \left[\frac{F_{0v,k}}{F_{0d,k}} \cdot f_{v,k}(t) \cdot \frac{q_{v,k}(t)}{v_{v,k}(t)}\right. \\
&\qquad \left. + \frac{F_{0d,k+1}}{F_{0d,k}} \cdot f_{d,k+1}(t) \cdot \frac{q_{d,k+1}(t)}{v_{d,k+1}(t)} - f_{d,k}(t) \cdot \frac{q_{d,k}(t)}{v_{d,k}(t)}\right], \\
t_{0d,k} \cdot \frac{dq_{d,k}(t)}{dt} &= \frac{F_{0v,k}}{F_{0d,k}} \cdot f_{v,k}(t) \cdot \frac{q_{v,k}(t)}{v_{v,k}(t)} \\
&\qquad + \frac{F_{0d,k+1}}{F_{0d,k}} \cdot f_{d,k+1}(t) \cdot \frac{q_{d,k+1}(t)}{v_{d,k+1}(t)} - f_{d,k}(t) \cdot \frac{q_{d,k}(t)}{v_{d,k}(t)}.
\end{aligned}
\tag{S1.12}
$$

Note that the description of baseline CBF and CBV, and their functional and anatomical dependencies across depths in the AV is provided below.

Next, the CBV change in AV compartment at specific depth $k$ is again given by difference between blood flows entering and leaving the comportment:

$$\frac{dV_{d,k}(t)}{dt} = F_{v,k}(t) + F_{d,k+1}(t) - F_{d,k}(t),$$
$$\frac{dV_{d,k}(t)/V_{0d,k}}{dt} = \frac{1}{V_{0d,k}} \cdot \left[\frac{F_{0v,k} \cdot F_{v,k}(t)}{F_{0v,k}} + \frac{F_{0d,k+1} \cdot F_{d,k+1}(t)}{F_{0d,k+1}} - \frac{F_{0d,k} \cdot F_{d,k}(t)}{F_{0d,k}}\right],$$
$$\frac{dv_{d,k}(t)}{dt} = \frac{F_{0d,k}}{V_{0d,k}} \cdot \left[\frac{F_{0v,k}}{F_{0d,k}} \cdot f_{v,k}(t) + \frac{F_{0v,k+1}}{F_{0d,k}} \cdot f_{d,k+1}(t) - f_{d,k}(t)\right], \tag{S1.13}$$

Using the same derivation steps as above, we arrive to the final form of depth-specific relative CBV changes in the AV:

$$t_{0d,k} \cdot \frac{dv_{d,k}(t)}{dt} = \frac{F_{0v,k}}{F_{0d,k}} \cdot f_{v,k}(t) + \frac{F_{0v,k+1}}{F_{0d,k}} \cdot f_{d,k+1}(t) - f_{d,k}(t). \tag{S1.14}$$

*Pial vein compartment:*

The PV is considered as vascular compartment at depth $k = 0$, which receives the output from AV. Within PV, the dynamic changes in dHb content are described by dHb flux entering from the top depth ($k = 1$) of AV and dHb flux leaving the PV compartment:

$$\frac{dQ_p(t)}{dt} = F_{d,1}(t) \cdot C_{d,1}(t) - F_p(t) \cdot C_p(t),$$
$$\frac{dq_p(t)}{dt} = \frac{F_{0d,1} \cdot C_{0d,1}}{Q_{0p}} \cdot f_{d,1}(t) \cdot \frac{C_{d,1}(t)}{C_{0d,1}} - \frac{F_{0p} \cdot C_{0p}}{Q_{0p}} \cdot f_p(t) \cdot \frac{C_p(t)}{C_{0p}} \tag{S1.15}$$

By assuming that the baseline CBF entering and leaving PV is the same (as no further vessels merging or branching is present at this level), $F_{0p} = F_{0d,1}$, and as above that the baseline dHb concentrations in venous vessels equal, $C_{0p} = C_{0d,1}$, the final form of relative changes in dHb content in the PV is given by:

$$t_{0p} \cdot \frac{dq_p(t)}{dt} = f_{d,1}(t) \cdot \frac{q_{d,1}(t)}{v_{d,1}(t)} - f_p(t) \cdot \frac{q_p(t)}{v_p(t)}, \tag{S1.16}$$

The relative CBV change in the PV is:

$$\frac{dV_p(t)}{dt} = F_{d,1}(t) - F_p(t),$$
$$\frac{V_{0p} \cdot dv_p(t)}{dt} = F_{0d,1} \cdot f_{d,1}(t) - F_{0p} \cdot f_p(t) \tag{S1.17}$$
$$t_{0p} \cdot \frac{dv_p(t)}{dt} = f_{d,1}(t) - f_p(t)$$

### 2. Derivation of laminar BOLD signal equation

The MR signal at rest (i.e. baseline) is a $CBV_0$ weighted sum of extravascular (EV) and intravascular (IV) signal components, distinguished with subscripts $E$ and $I$, respectively:

$$S_{0,k} = \left(1 - \bar{V}_{0v,k} - \bar{V}_{0d,k}\right) \cdot S_{0E_k} + \bar{V}_{0v,k} \cdot S_{0I_{v,k}} + \bar{V}_{0d,k} \cdot S_{0I_{d,k}}, \tag{S1.18}$$

where $\bar{V}_{0i,k}$ is the fractional $CBV_0$ of blood with respect to GM tissue, specific for $k$-th cortical depth and vascular compartment, i.e. in contrast to $V_{0i,k}$ used in the dynamic equations above, the $CBV_0$ values are further divided by the amount of surrounding tissue. For conversion between $V_{0i,k}$ and $\bar{V}_{0i,k}$ see below section Baseline CBV and CBF. The extravascular and intravascular signals are modeled as:

$$\begin{aligned} S_{0E_k} &= \rho_E \cdot M_z \cdot e^{-TE \cdot R_{2E_k}^*}, \\ S_{0I_{v,k}} &= \rho_{I_v} \cdot M_z \cdot e^{-TE \cdot R_{2I_{v,k}}^*}, \\ S_{0I_{d,k}} &= \rho_{I_d} \cdot M_z \cdot e^{-TE \cdot R_{2I_{d,k}}^*}, \end{aligned} \tag{S1.19}$$

where $\rho_E$ and $\rho_{I_i}$ are water proton densities in the GM tissue and blood, respectively. $\rho_{I_i}$ can be calculated from $\rho_{I_i} = 0.95 - 0.22 \cdot Hct_i$ (Lu et al., 2002), where $Hct_i$ is the fractional hematocrit in the blood. $TE$ is the echo time and $R_{2E_k}^*$ and $R_{2I_{i,k}}^*$ are the baseline EV and IV (apparent) transverse relaxation rates. The effect of longitudinal magnetization $M_z$ (which could potentially reflect laminar variation in longitudinal relaxation time $T_1$ and contribution of inflow effect) is not modeled below (i.e. set to one).

With activation, the EV and IV transverse relaxation rates are altered by additive amounts $\Delta R_{2E_{i,k}}^*$ and $\Delta R_{2I_{i,k}}^*$, and the CBV changes to new values $\bar{V}_{i,k} = v_{i,k} \cdot \bar{V}_{0i,k}$. The MR signal with activation is then:

$$\begin{aligned} S_k = \left(1 - v_{v,k} \cdot \bar{V}_{0v,k} - v_{d,k} \cdot \bar{V}_{0d,k}\right) \cdot S_{0E_k} \cdot e^{-TE \cdot \left(\Delta R_{2E_{v,k}}^* + \Delta R_{2E_{d,k}}^*\right)} + v_{v,k} \cdot \bar{V}_{0v,k} \\ \cdot S_{0I_{v,k}} \cdot e^{-TE \cdot \Delta R_{2I_{v,k}}^*} + v_{d,k} \cdot \bar{V}_{0d,k} \cdot S_{0I_{d,k}} \cdot e^{-TE \cdot \Delta R_{2I_{d,k}}^*}. \end{aligned} \tag{S1.20}$$

One can see that for given $TE$ and $CBV_0$ distribution $\bar{V}_{i,k}$, the signal $S_k$ is a function of $\Delta R_{2E_{i,k}}^*$, $\Delta R_{2I_{i,k}}^*$ and $v_{i,k}$. We now approximate the change in MR signal from baseline to activation, $\Delta S_k = S_k - S_{0,k}$, using the first-order Taylor series expansion around equilibrium points $\Delta R_{2E_{i,k}}^* = 0, \Delta R_{2I_{i,k}}^* = 0$ and $v_{i,k} = 1$:

$$
\begin{aligned}
\Delta S_k &= S_k - S_{0,k} \\
&= (v_{v,k} - 1)\frac{\partial S_k}{\partial v_{v,k}} + (v_{d,k} - 1)\frac{\partial S_k}{\partial v_{d,k}} + \Delta R^*_{2E_{v,k}}\frac{\partial S_k}{\partial \Delta R^*_{2E_{v,k}}} \\
&+ \Delta R^*_{2E_{d,k}}\frac{\partial S_k}{\partial \Delta R^*_{2E_{d,k}}} + \Delta R^*_{2I_{v,k}}\frac{\partial S_k}{\partial \Delta R^*_{2I_{v,k}}} + \Delta R^*_{2I_{d,k}}\frac{\partial S_k}{\partial \Delta R^*_{2I_{d,k}}} \\
&= (v_{v,k} - 1)\cdot \bar{V}_{0v,k}\cdot\left(S_{0I_{v,k}} - S_{0E_k}\right) + (v_{d,k} - 1)\cdot \bar{V}_{0d,k}\cdot\left(S_{0I_{d,k}} - S_{0E_k}\right) \\
&\qquad - \left(1 - \bar{V}_{0v,k} - \bar{V}_{0d,k}\right)\cdot S_{0E_k}\cdot\left(\Delta R^*_{2E_{v,k}} + \Delta R^*_{2E_{d,k}}\right)\cdot TE \\
&- \bar{V}_{0v,k}\cdot S_{0I_{v,k}}\cdot \Delta R^*_{2I_{v,k}}\cdot TE - \bar{V}_{0d,k}\cdot S_{0I_{d,k}}\cdot \Delta R^*_{2I_{d,k}}\cdot TE.
\end{aligned}
\tag{S1.21}
$$

Expressing the MR signal as a fractional signal change (i.e. dividing by $S_{0,k}$) yields the BOLD signal equation:

$$
\begin{aligned}
\frac{\Delta S_k}{S_{0k}} = &\frac{1}{\left(1 - \bar{V}_{0v,k} - \bar{V}_{0d,k}\right) + \varepsilon_{d,k}\cdot \bar{V}_{0v,k} + \varepsilon_{d,k}\cdot \bar{V}_{0d,k}}\Big[\bar{V}_{0v,k}\cdot (v_{v,k} - 1) \\
&\cdot (\varepsilon_{v,k} - 1) + \bar{V}_{0d,k}\cdot (v_{d,k} - 1)\cdot (\varepsilon_{d,k} - 1) - \left(1 - \bar{V}_{0v,k} - \bar{V}_{0d,k}\right) \\
&\cdot \left(\Delta R^*_{2E_{v,k}} + \Delta R^*_{2E_{d,k}}\right)\cdot TE - \varepsilon_{v,k}\cdot \bar{V}_{0v,k}\cdot \Delta R^*_{2I_{v,k}}\cdot TE - \varepsilon_{d,k}\cdot \bar{V}_{0d,k} \\
&\cdot \Delta R^*_{2I_{d,k}}\cdot TE\Big],
\end{aligned}
\tag{S1.22}
$$

where we used substitution $\varepsilon_{i,k} = S_{0I_{i,k}}/S_{0E,k}$, which refers to the ratio of baseline IV-to-EV signals.

Next, by using results of (Ogawa et al., 1993), we express changes in transverse relaxation rates in terms of physical and physiological variables (as described in (Obata et al., 2004) or (Stephan et al., 2007)). The cortical depth-dependent changes in EV $R^*_{2E}$ components are given by:

$$
\begin{aligned}
\Delta R^*_{2E_{i,k}} &= 4.3\cdot \vartheta_{0i}\cdot\left[\bar{V}_{i,k}\cdot\left(1 - Y_{i,k}\right) - \bar{V}_{0i,k}\cdot (1 - Y_0)\right] \\
&= 4.3\cdot \vartheta_{0i}\cdot \bar{V}_{0i,k}\cdot (1 - Y_0)\cdot\left[\frac{\bar{V}_{i,k}\cdot\left(1 - Y_{i,k}\right)}{\bar{V}_{0i,k}\cdot (1 - Y_0)} - 1\right] \\
&= 4.3\cdot \vartheta_{0i}\cdot \bar{V}_{0i,k}\cdot E_0\cdot\left(q_{i,k} - 1\right).
\end{aligned}
\tag{S1.23}
$$

Note this expression is valid for venules and large veins (including AV and PV) (Ogawa et al., 1993). $(1 - Y_{i,k})$ is the deoxygenation of the blood in specific vascular compartment and cortical depth; i.e. it is the oxygen extraction fraction, $E_{i,k}$, and $(1 - Y_0) = E_0$. Note that we dropped the depth-specific and vascular-compartment-specific subscripts in the baseline oxygen extraction fraction because it is assumed constant across all venous vessels. Here $\vartheta_{0i} = \Delta\chi_0\cdot Hct_i\cdot\gamma\cdot B_0$, where $\Delta\chi_0$ is the susceptibility difference between of fully oxygenated and deoxygenated blood, $B_0$ refers to magnetic field strength, and $\gamma$ is the gyromagnetic ratio of water protons (see Table 2 for further details). Above we also used substitution:

$$
q_{i,k} = \frac{Q_{i,k}}{Q_{0i,k}} = \frac{\bar{V}_{i,k}\cdot\left(1 - Y_{i,k}\right)}{\bar{V}_{0i,k}\cdot (1 - Y_0)}.
\tag{S1.24}
$$

The cortical depth-dependent changes in IV $R^*_2$ components are given by (Obata et al., 2004):

$$
\begin{aligned}
\Delta R^*_{2I_{i,k}} &= r_{0i} \cdot \left[\left(1 - Y_{i,k}\right) - \left(1 - Y_0\right)\right] \\
&= r_{0i} \cdot \left(1 - Y_0\right) \cdot \left[\frac{\left(1 - Y_{i,k}\right)}{\left(1 - Y_0\right)} - 1\right] \\
&= r_{0i} \cdot E_0 \cdot \left(\frac{q_{i,k}}{v_{i,k}} - 1\right),
\end{aligned} \tag{S1.25}
$$

where $r_{0i}$ is the slope of changes in IV signal relaxation rate with respect to expected changes in extraction fraction $(1 - Y)$ and we used substitution, demonstrating that relative dHb concentration equals relative oxygen extraction fraction:

$$
\frac{q_{i,k}}{v_{i,k}} = \frac{Q_{i,k}}{Q_{0i,k}} \cdot \frac{\bar{V}_{0i,k}}{\bar{V}_{i,k}} = \frac{\left(1 - Y_{i,k}\right)}{\left(1 - Y_0\right)}. \tag{S1.26}
$$

By inserting S1.23 and S1.25 to the BOLD signal equation S1.22 gives:

$$
\begin{aligned}
\frac{\Delta S_k}{S_{0,k}} = H_{0,k} \Big[ & \bar{V}_{0v,k} \cdot \left(v_{v,k} - 1\right) \cdot \left(\varepsilon_v - 1\right) + \bar{V}_{0d,k} \cdot \left(v_{d,k} - 1\right) \cdot \left(\varepsilon_d - 1\right) \\
& - \left(1 - \bar{V}_{0v,k} - \bar{V}_{0d,k}\right) \cdot 4.3 \cdot E_0 \cdot TE \\
& \cdot \left(\vartheta_{0v} \cdot \bar{V}_{0v,k} \cdot \left(q_{v,k} - 1\right) + \vartheta_{0d} \cdot \bar{V}_{0d,k} \cdot \left(q_{d,k} - 1\right)\right) - \varepsilon_{v,k} \cdot \bar{V}_{0v,k} \\
& \cdot r_{0v} \cdot E_0 \cdot \left(\frac{q_{v,k}}{v_{v,k}} - 1\right) \cdot TE - \varepsilon_{d,k} \cdot \bar{V}_{0d,k} \cdot r_{0d} \cdot E_0 \cdot \left(\frac{q_{d,k}}{v_{d,k}} - 1\right) \\
& \cdot TE \Big],
\end{aligned} \tag{S1.27}
$$

where we used substitution $H_{0,k} = 1/\left(1 - \bar{V}_{0v,k} - \bar{V}_{0d,k} + \varepsilon_{v,k} \cdot \bar{V}_{0v,k} + \varepsilon_{d,k} \cdot \bar{V}_{0d,k}\right)$.

By grouping constants together with respect to relative EV, IV and CBV, across different vascular compartments:

$$
\begin{aligned}
c_{1i} &= 4.3 \cdot \Delta\chi_0 \cdot Hct_i \cdot \gamma \cdot B_0 \cdot E_0 \cdot TE, \\
c_{2i,k} &= \varepsilon_{i,k} \cdot r_{0i} \cdot E_0 \cdot TE, \\
c_{3i,k} &= 1 - \varepsilon_{i,k},
\end{aligned} \tag{S1.28}
$$

the final form of the laminar BOLD signal equation yields the following form:

$$
\begin{aligned}
\frac{\Delta S_k(t)}{S_k} = H_{0,k} \cdot \Bigg[ & \left(1 - \sum_i \bar{V}_{0i,k}\right) \cdot \sum_i c_{1i} \cdot \bar{V}_{0i,k} \cdot \left(1 - q_{i,k}(t)\right) \\
& + \sum_i c_{2i,k} \cdot \bar{V}_{0i,k} \cdot \left(1 - \frac{q_{i,k}(t)}{v_{i,k}(t)}\right) + \sum_i c_{3i,k} \cdot \bar{V}_{0i,k} \cdot \left(1 - v_{i,k}(t)\right) \Bigg]
\end{aligned} \tag{S1.29}
$$

*BOLD signal equation for PV:*

PV is represented by a single vascular comportment. Derivation of its BOLD signal equation follows exactly the same steps as shown above. Starting from MRI signal during rest, the EV and IV signal components are weighted by CBV₀, $\bar{V}_{0p}$, which is defined independently of the CBV₀ within GM tissue:

$$S_{0p} = \left(1 - \bar{V}_{0p}\right) \cdot S_{0E_p} + \bar{V}_{0p} \cdot S_{0I_p}. \tag{S1.30}$$

Baseline EV and IV signal components are defined as in (S1.19) but see also Table 2 for details about parameterization. With activation:

$$S_p = \left(1 - v_p \cdot \bar{V}_{0p}\right) \cdot S_{0E_p} \cdot e^{-TE \cdot \Delta R^*_{2E_p}} + v_p \cdot \bar{V}_{0p} \cdot S_{0I_p} \cdot e^{-TE \cdot \Delta R^*_{2I_p}}. \tag{S1.31}$$

Approximation of the change in MR signal from baseline to activation, $\Delta S_p = S_p - S_{0p}$, using the first-order Taylor series expansion around equilibrium points $\Delta R^*_{2E_p} = 0, \Delta R^*_{2I_p} = 0$ and $v_p = 1$, gives:

$$\begin{aligned} \Delta S_p &= S_p - S_{0p} = \\ &= \left(v_p - 1\right)\frac{\partial S_p}{\partial v_p} + \Delta R^*_{2E_p}\frac{\partial S_p}{\partial \Delta R^*_{2E_p}} + \Delta R^*_{2I_p}\frac{\partial S_p}{\partial \Delta R^*_{2I_p}} \\ &= \left(v_p - 1\right) \cdot \bar{V}_{0p} \cdot \left(S_{0I_p} - S_{0E_p}\right) - \left(1 - \bar{V}_{0p}\right) \cdot S_{0E_p} \cdot \Delta R^*_{2E_p} \cdot TE - \bar{V}_{0p} \cdot S_{0I_p} \\ &\quad \cdot \Delta R^*_{2I_p} \cdot TE \end{aligned} \tag{S1.32}$$

Expressing the MR signal as a fractional signal change (i.e. dividing by $S_{0p}$) yields the BOLD signal equation for PV:

$$\begin{aligned} \frac{\Delta S_p}{S_{0p}} &= \frac{S_{0E_p}}{\left(1 - \bar{V}_{0p}\right) \cdot S_{0E_p} + \bar{V}_{0p} \cdot S_{0I_p}}\left[\left(v_p - 1\right) \cdot \bar{V}_{0p} \cdot \left(\frac{S_{0I_p}}{S_{0E_p}} - 1\right) - \left(1 - \bar{V}_{0p}\right)\right. \\ &\quad \left. \cdot \Delta R^*_{2E_p} \cdot TE - \bar{V}_{0p} \cdot \frac{S_{0I_p}}{S_{0E_p}} \cdot \Delta R^*_{2I_p} \cdot TE\right] \\ &= \frac{\bar{V}_{0p}}{\left(1 - \bar{V}_{0p}\right) + \varepsilon_p \cdot \bar{V}_{0p}}\left[\left(1 - \bar{V}_{0p}\right) \cdot c_{1p} \cdot \left(1 - q_p\right) + c_{2p} \cdot \left(1 - \frac{q_p}{v_p}\right) + c_{3p}\right. \\ &\quad \left. \cdot \left(1 - v_p\right)\right], \end{aligned} \tag{S1.33}$$

where we assume that changes in EV and IV transversal relaxation rates, $\Delta R^*_{2E_p}$ and $\Delta R^*_{2I_p}$, follow the same expressions as in (S1.23 and S1.25), with PV-specific parameters defined in Table 2.

### 3. Baseline CBV and CBF

For dynamic equations (e.g. S1.4 and S1.9), we consider the total amount of CBV₀ ($V_0$) in the units of mL, which is divided between venules and AV comportments and a specific number of depths:

$$V_{0i,k} = \omega_i \cdot x_{i,k} \cdot V_0 \tag{S1.34}$$

Here, $\omega_i$ refers to fractions of venules and AV, and $x_{i,k}$ are fractions describing blood volume distribution between different depths, which can be different for MV and AV. In principle, one is allowed to freely define the $CBV_0$ distribution in both MV and AV. However, in order to make the description of the model easier and independent of number of depths, we chose to parameterize the increase (or even decrease) of $CBV_0$ towards the surface either in MV or AV by assumining a linear function:

$$\tilde{x}_{i,k} = s_{0,i} + s_i \cdot (1 - l_k) \cdot 100, \tag{S1.35}$$

where $l_k$ refers to the normalized distance (i.e. between 0 and 1) for $k$-th depth with respect to the CSF surface. $s_{0,i}$ refers to a positive offset that ensures non-zero blood volume in each depth (in our simulations we asummed $s_{0,i} = 10$) and $s_i$ is a slope linear increase (i.e. positive constant) towards the surface. The normalized fractions $x_{i,k}$ in (S1.35) are then obtained by dividing $\tilde{x}_{i,k}$ with its total sum across all cortical depths. Note that in the paper, we formalized the increase of $CBV_0$ towards the surface using ratio between superficial and deepest depth, but this definition is not independent of number of depths. As mentioned above, for PV we define total $CBV_0$, $V_{0p}$, independently of total amount of $CBV_0$ for MV and AV.

Further, in the laminar BOLD signal equation (S1.22) the baseline CBV is represented as a fraction with respect to surrounding GM tissue (i.e. typically per 100 g of tissue). By assuming that the amount of gray matter tissue (GM) is divided into $K$ equivolume depths (i.e. 100 g$/K$), the depth-specific blood volume *fraction* of venules and AV with respect to tissue is obtained as:

$$\bar{V}_{0i,k} = V_{0i,k} \cdot K/100. \tag{S1.36}$$

Next, the $CBF_0$ is expressed in units of mL of blood per sec. For a given distribution of $CBV_0$ in the MV, we can calculate depth-specific $CBF_0$, by assuming specific transit time (in units of sec) through MV, $F_{0v,k} = V_{0v,k}/t_{0v,k}$. Note that in all presented simulations, we considered constant transit time through MV across depths. However, it is also possible to consider that it varies across cortical depths, simply by defining $t_{0v,k}$ as depth-specific. Additionally, we assume that the the transit time through MV does not change with different number of depths. However, the depth-specific flow in MV, $F_{0v,k}$, does change, because the $CBV_0$ distribution changes (but the total flow $\sum_{k=1}^{K} F_{0v,k}$ remains the same). Considering the AV, $CBF_0$ follows the mass conservation law (i.e. continuity equation in particular). That is, the $CBF_0$ in $k$-th depth is a sum of blood flow leaving the MV of that depth, and the blood flow from the lower depth (i.e. $k+1$) of the AV:

$$F_{0d,k} = F_{0v,k} + F_{0d,k+1}. \quad \text{(S1.37)}$$

For the lowest depth ($k = K$), $CBF_0$ in the AV equals the blood flow from MV of the same depth, $F_{0d,K} = F_{0v,K}$. Thus, one can also see that the $CBF_0$ in specific depth is dependent on the number of selected depth, but the outflow from the AV compartment in the superficial depth ($k = 1$) will be independent of the number of depth, i.e. equal total flow in the MV $F_{0d,1} = \sum_{k=1}^{K} F_{0v,k}$. Further, $CBF_0$ in the PV equals the total $CBF_0$ leaving the AVs, $F_{0d,1} = F_{0p}$. In the Table S1, we summarize, which physiological baseline parameters are affected by different number of depths and which are not. In general, all parameters in the model (i.e. describing baseline conditions, steady-state and dynamic relationships between physiological variables), can be defined as depth-specific, with $E_0$ as the only exception. The dynamic mass balance equations were derived under the assumption of $E_{0i,k}$ (or $C_{0i,k}$) being constant everywhere in across venous vasculature. That is, a new and more complex dynamic equtions would have to be derived in order to allow for some variability in $E_{0i,k}$ across depths and venous compartments. Nevertheless, it is expected that small variation in $E_{0i,k}$ across depth would have minimal effect on variability in laminar BOLD response.

Note that the CBF, as defined in our model, is closely related to blood perfusion measured arterial spin labeling (ASL) technique (Liu and Brown, 2007), commonly expressed in units of mL of blood per min per 100 g of tissue. This is especially the case if we talk about the input of our model representing the blood flow leaving arterioles. Then, within dynamic equations and without loss of generality, it is also possible to express $CBF_0$ as a flow rate (e.g. perfusion 75 mL/min/100g in MV equals to flow rate 0.0125 $s^{-1}$) and $CBV_0$ in fractions (e.g. 0.0125), as long as the resulting transit time is in units of sec. In the Supplementary Material 2, we provide an example how considering different number of depths is reflected in the baseline parameters and laminar BOLD response.

**Table S1**. Summary of physiological baseline parameters, which are number of depths depdendent and independent.

| Parameter | No. of depths dependent | No. of depths independent |
|---|---|---|
| $V_{0v,k}$ | ● | - |
| $V_{0d,k}$ | ● | - |
| $F_{0v,k}$ | ● | - |
| $F_{0d,k}$ | ● | - |
| $t_{0v,k}$ | - | ● |
| $t_{0d,k}$ | ● | - |
| $E_{0v,k}$ | - | ● |
| $E_{0d,k}$ | - | ● |
