## Supplementary Material 2 (Additional simulations and results) for "A dynamical model of the laminar BOLD response"


### 1. Additional factors affecting the LBR

Figure S2.1A shows that lowering CBV change in the MV has mostly a simple additive effect on LBR increase towards the surface. On the other hand, lowering CBV change in the AV has mostly simple scaling (i.e. multiplicative) effect on LBR (see Figure S2.1B). Similar scaling

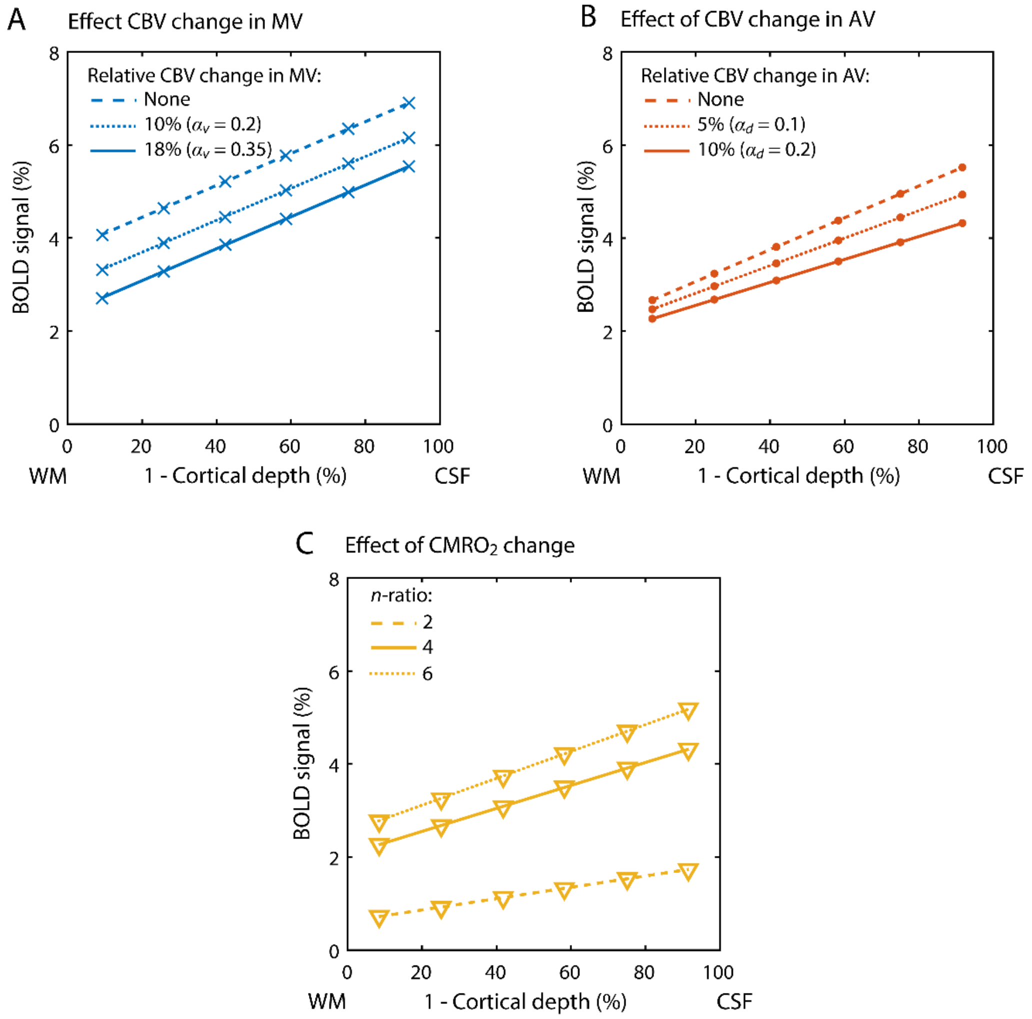


**Figure S2.1.** Simulated LBRs during steady-state depicted as function of (1 - normalized cortical depth) between WM and CSF boundaries. (A) LBR dependence on relative CBV change in the MV and (B) in the AV. (C) LBR dependence on relative $CMRO_2$ change (parameterized with $n$-ratio with respect to relative CBF change).

effect can be also achieved by increasing the ratio between relative CBF and $CMRO_2$ (see Figure S2.1C). Here, in combination with CBV change in both MV and AV, lower $n = 2$ results in relatively low amplitude LBR (dashed line).

### 2. Dependence of LBR and baseline parameter on number of depths

If one wants to compare how considering different number of depths affects LBR and underlying baseline parameters, it is important to realize that the underlying physiology does not change (only our way we 'look' at it). That is, given the mass conservation law, the total baseline physiological parameters must remain the same independent of the number of depths. Here, we consider our default steady-state scenario and evaluate it for 3 and 6 depths. In Figure S2.2A, we see that the baseline CBV (in mL) per depth in the AV reduced with 6 depths (solid line) compared to 3 depths (dashed line) but the total amount of baseline CBV in AV is the same. Note that if we depict the baseline CBV in fractions with respect to the GM tissue (Figure S2.2B), we see that with the different number of depths we only sample at different location along the same line. Laminar baseline CBV distribution in fractions can be linked to vascular

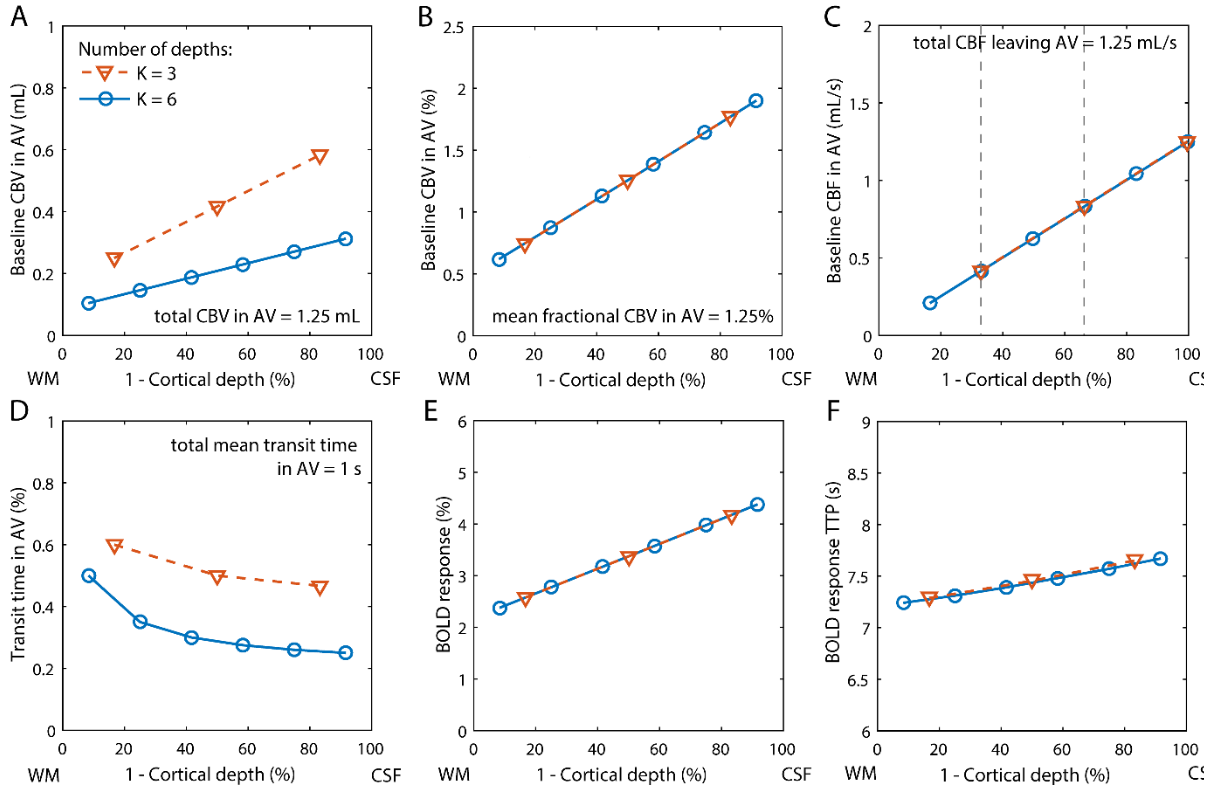


**Figure S2.2.** Comparison between modeling 3 and 6 cortical depths of LBR. (A) Baseline CBV in expressed in mL. (B) Baseline CBV in AV expressed as a faction in percent with respect to the surrounding tissue. (C) Baseline CBF (outflow) within AV expressed in mL/s, depicted on the edge of physical border between different depths. (D) Transit time through compartments of AV. (E) LBR profiles. (F) TTP of LBR with respect to stimulus onset.

density, which is reported in some animal studies (e.g.(Schmid et al., 2017)). Next, in Figure S2.2C, we see how the baseline CBF as an outflow (in mL per s) leaving compartments in AV accumulates towards the surface. Note that the baseline CBF becomes twice smaller per depth in the MV (assuming homogenous distribution), which is also noticeable in the lowest depth of AV. One can notice that the outflows displayed as a function of cortical depth for 3 and 6 depths lay on the same line, and the total outflow leaving the AV is independent of the number of depths. Here we illustrate that outflows from AV compartments are leaving segments of tissues in different depths, and therefore, they are aligned to upper edge of each depth rather than its center. As a result of laminar baseline CBV (in mL) and CBF (in mL per sec) on number of depths, with increasing number of depths the transit times through AV compartments get shorter (see Figure S2.2D), but again the total *mean* transit time in the AV remains the same (defined as a mean over a cumulative sum of depth-specific transit times in the AV, starting from the depth closest to the surface). Next, as for the laminar distribution of baseline CBV fractions, the LBR profiles for 3 and 6 depths lay also along the same line (see Figure S2.2E), and this is almost the case also for depth-dependence of LBR-TTP (see Figure S2.2F).

### 3. Effect of variable laminar CBF response on LBR

In order to further explore the effect of variable laminar CBF response on LBR, we consider a scenario where there is an increase of relative CBF (60%) only in the lower and upper depths but no increase in the middle depths (see Figure S2.3A). Further, we have assumed a case with and without baseline CBV increasing towards the surface in the AV. All other model parameters were the same as in our default steady-state scenario. The resulting LBRs for both cases are shown in Figure S2.3B. With the baseline CBV increasing towards the surface in AV (solid line), we see that although the LBR drops in the middle depths, there is still a general tendency of LBR to increase towards the surface (i.e. we observe larger signal amplitude in the upper depths compared to the lower depths). This scenario captures effect of both baseline CBV increase and dHb drainage towards the surface in the AV. On the other hand, without baseline CBV increasing towards the surface (i.e. being constant across depths), we observe a lower amplitude of LBR in the upper depth compared to the lower depths (dashed line). This is because for constant CBV in the AV (physiologically less supported scenario), the transit time in the upper depth is much shorter (about six times in this case) compared to the lowest depth. This shows again that only dHb drainage to upper depths is (physiologically) not enough to explain the typical increase of LBR towards the surface.

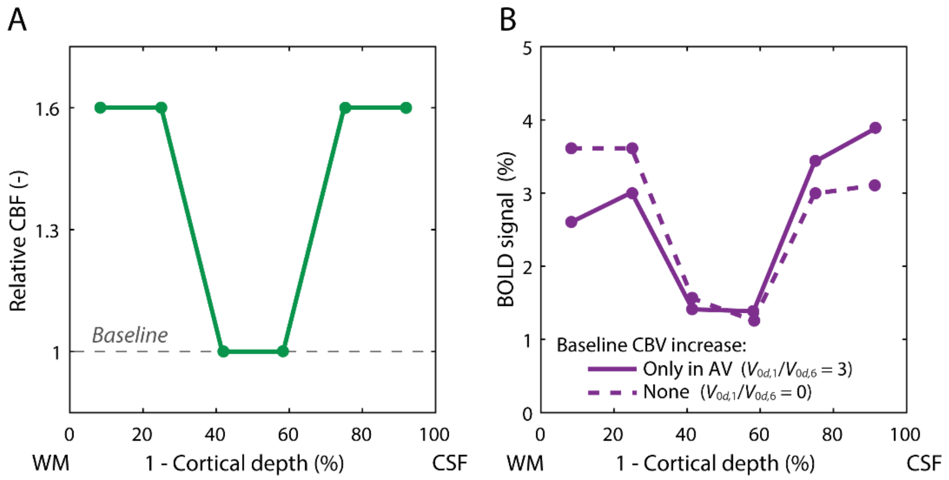


**Figure S2.3** (A) Laminar profile of relative CBF change. (B) Resulting LBRs for baseline CBV increasing towards the surface (solid line) and being constant across depths (dashed line) in the AV.

### 4. Additional factor affecting LBR transients

Changes in $CBV_0$ ratio between the MV and AV can significantly modulate TTP and TTU between depths (see Figure S2.4A). For example, increasing the proportion of baseline $CBV_0$ in the AV with respect to the microvasculature results in more pronounced delays of response transients between depths (i.e. the difference of TTP and TTU between lower and upper depths become larger). Next, increasing the transit time through MV has mostly simple scaling effect on TTP and TTU across depths (see Figure S2.4B). Similar effect is observed also on amplitude increase towards the surface during PSU (Figure S2.4B, third column).

.

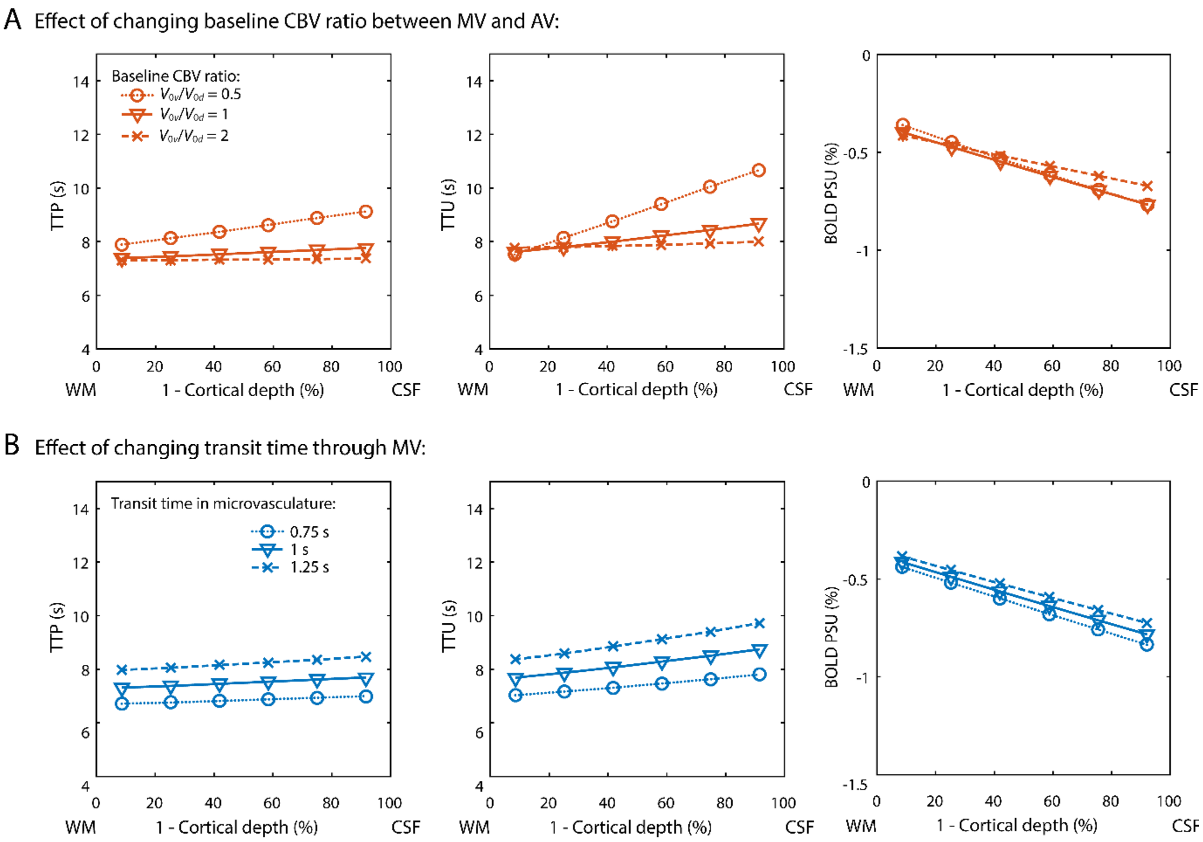


**Figure S2.4** Simulation results demonstrating the effect of $CBV_0$ ratio between MV and AV (A) and transit-time in MV (B) on laminar dependence of temporal BOLD response features such as TTP (first column), TTU (second column) and amplitude of PSU (third column). Note that the TTU is calculated with respect to the end of stimulus.
